# The fruit fly, *Drosophila melanogaster*, as a micro-robotics platform

**DOI:** 10.1101/2024.05.24.595748

**Authors:** Kenichi Iwasaki, Charles Neuhauser, Chris Stokes, Aleksandr Rayshubskiy

## Abstract

Engineering small autonomous agents capable of operating in the microscale environment remains a key challenge, with current systems still evolving. Our study explores the fruit fly, *Drosophila melanogaster*, a classic model system in biology and a species adept at microscale interaction, as a biological platform for micro-robotics. Initially, we focus on remotely directing the walking paths of fruit flies in an experimental arena. We accomplish this through two distinct approaches: harnessing the fruit flies’ opto-motor response and optogenetic modulation of its olfactory system. These techniques facilitate reliable and repeated guidance of flies between arbitrary spatial locations. We guide flies along predetermined trajectories, enabling them to scribe patterns resembling textual characters through their locomotion. We enhance olfactory-guided navigation through additional optogenetic activation of positive valence mushroom body output neurons. We extend this control to collective behaviors in shared spaces and navigation through constrained maze-like environments. We further use our guidance technique to enable flies to carry a load across designated points in space, establishing the upper bound on their weight carrying capabilities. Additionally, we demonstrate that visual guidance can facilitate novel interactions between flies and objects, showing that flies can consistently relocate a small spherical object over significant distances. Beyond expanding tools available for micro-robotics, these novel behavioral contexts can provide insights into the neurological basis of behavior in fruit flies.

## Main Text

Micro-robots hold promise in their ability to perform collective tasks and to navigate in spaces that are otherwise inaccessible to larger robots or humans. Their applications include environmental surveillance and precision agriculture, where they could discreetly monitor air quality, aid in pollination, or gather critical data with minimal ecological impact. In search and rescue operations in disaster-stricken areas, micro-robotic swarms could traverse rubble and tight quarters to locate survivors or assess structural damage. There is a concerted effort to engineer micro-robots with proficient ambulatory (*1–3*) and aerial (*4–6*) capabilities, often taking inspiration from insects (*7*). Recent advancements in micro-robotics have been significant, yet they are hindered by several limitations. The current computational frameworks governing their locomotion and navigational abilities are constrained due to the limited onboard space needed for sophisticated control systems that facilitate complex tasks. Moreover, the assembly and manufacturing processes for these devices is complex and laborious, demanding sophisticated techniques and designs tailored for microscale production (*8, 9*).

In this study, we propose the use of a small (approximately 1.5 mm wide x 2.5 mm long) biological organism, the fruit fly, *Drosophila melanogaster* as a fundamental unit of a micro-robotics platform. For millions of years, fruit flies have evolved a set of abilities to maneuver seamlessly in the natural environment (*10*). Fruit flies are also one of the oldest genetics model systems in biology, having yielded genetic tools that today allow us to precisely target for manipulation many neurons in their brain – leading to remote (optogenetic) neural control at the level of a single cell type (*11*). In addition, there is extensive and rapidly expanding knowledge on the neural circuits that control sensory processing (*12–15*), navigation (*16*), motor control (*17–19*) and high level behaviors such as courtship (*20*) and fighting (*21*), feeding (*22*), egg-laying (*23*) and grooming (*19, 24*). Many of these behaviors can already be triggered with optogenetic activation. Our ability to control various behaviors has the potential to further improve with the use of a detailed neural circuit diagram (*25*), enabling further insight into neural circuits that regulates behavior in fruit flies. Recently new efforts have emerged to perform whole body simulations of the fruit fly, using anatomically-detailed biomechanical whole-body model equipped with neural controllers for walking, grooming (*26*) and flight (*27*). Furthermore, fruit flies can be bred reliably and economically at scale, avoiding the complexities of robotic fabrication. The extensive array of tools and research insights available today presents an unparalleled opportunity to re-evaluate the fruit fly, a classical biological model system, within the framework of a micro-robotics.

Here, we present two methods, each with distinct advantages, to direct the walking path of unrestrained fruit flies, leveraging two extensively studied behavioral responses to visual and olfactory stimuli. The visual approach depends on moving visual stimuli eliciting an opto-motor turning response (*28, 29*). In this method, we project a rotating pinwheel of alternating blue and black stripes over the fly and control the flies’ walking direction by rotating the pinwheel as the fly walks towards precise spatial goals. Using visual guidance, flies can follow the ‘steering’ command 94% of the time. The olfactory approach depends on the ability of flies to turn in response to an asymmetrical olfactory gradient, a behavior called osmotropotaxis (*30*). We developed a method to drive turning by asymmetrically stimulating the olfactory system during free walking. Using this method we show that we can guide flies to follow our ‘steering’ command to an arbitrary spatial position in the arena about 80% of the time. We further improve on this method by additionally activating several classes of Mushroom Body Output Neurons (MBONs) that have previously been shown to be involved in positive behavioral valance in the context of olfaction. We compare both of these methods, and demonstrate: fine-tuned navigational control with temporal persistence, enhanced olfactory navigation using genetic manipulations, and ability of flies to trace intricate spatial patterns akin to ’writing text’. We have coordinated simultaneous ’writing’ by multiple flies, steered them through constrained maze-like environments, and directed them to transport cargo over substantial distances, carrying weights almost equal to their own body weight for several hundred meters. Furthermore, we demonstrate that visual guidance can facilitate novel interactions between flies and objects, enabling a fly to relocate a 10mg ball over tens of centimeters.

## Results

### Visual-based remote control of the fly’s walking path

Fruit flies’ walking direction can be guided by moving visual stimuli via the optomotor response (*31*). The optomotor response in fruit flies is a well-studied phenomenon demonstrating their ability to stabilize their walking path in response to visual stimuli. This reflexive behavior is elicited when moving patterns are presented to the visual field of the fly. Essentially, the fly attempts to maintain a steady course by adjusting its flight or walking direction in response to perceived motion. For instance, if a pattern moves from left to right in the fly’s field of view, the fly will turn to the right, effectively trying to follow the motion (*28, 32*).

We set up an environment where we track flies using a camera and project a striped, black and blue circular pinwheel pattern centered at the fly (Figure 1A). When the pinwheel rotates clockwise, we expect to elicit a right turn in the fly, and when it rotates counterclockwise we expect to elicit a left turn (Figure 1Bi). Using this approach, we guide flies to walk in a line, back and forth between two points in space that are 13cm apart. Examples of individual running trajectories between regions A and B are shown in Figure 1C, displaying clear turning to clockwise and counterclockwise rotations of the pinwheel. For most trials, flies followed our guidance, staying roughly within the linear area between regions A and B (Figure 1D). Fidelity scores for visually-guided flies were typically ∼94% for BerlinK females (Figure 1F), with robust turning responses to clockwise and counterclockwise pinwheel rotations (Figure 1E). In addition to wildtype BerlinK flies, we examined the ability of a common ‘Rubin collection’ genetic background (*33*) to perform this task (Figure 1F,G). Flies can perform hundreds of successful runs to goal (Figure 1G) for 10s of hours (Supplementary Figure 4D) and many flies display high level of pinwheel-following behavior for hundreds of consecutive successful runs between regions A and B (Figure 1H).

**Figure 1:**
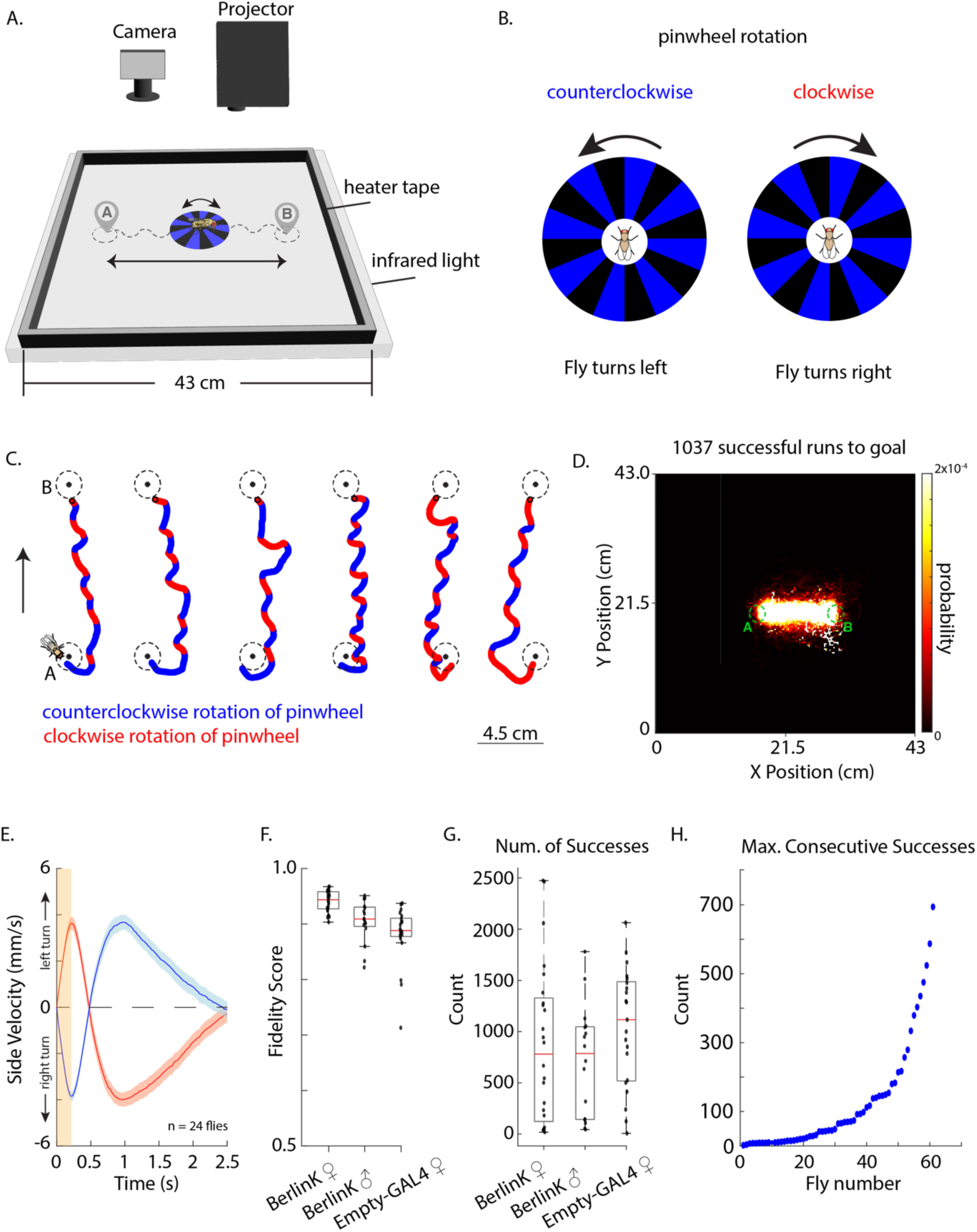
Visual based directed turning during free walking. A. Setup depicting projector assisted, pinwheel-based visual guidance in a large experimental arena. In this example, the fly is guided back and forth between spatial locations A and B, while the pinwheel is rotating clockwise or counterclockwise to guide the flies’ walking direction. B. Schematic depicting the stimulation protocol. Clockwise rotations of the pinwheel drive right turning and counterclockwise rotations drive left turning. C. Example fly running trajectories between point A to point B. Direction of pinwheel rotation is marked with red for clockwise and blue for counterclockwise. Flies’ orientation with respect to the running trajectory is noted by the fly cartoon on the left. D. An example fly’s probability density histogram is shown of the flies’ occupancy in the entire experimental arena over the course of the entire experiment. The fly was guided to run back and forth between regions A and B, completing 1037 successful trials, as defined by reaching a goal within 60 seconds. E. Average and SEM of side velocity aligned to the onset of pinwheel rotation at t = 0s (n = 24 flies, BerlinK females). Note, there is a delay of approximately 200ms before the fly begins executing an expected turn, because the fly is already executing a turn when the rotation direction switches (represented by an orange bar) and the fly needs some time to switch its behavioral turn to the opposite direction. This delay corresponds to the movement of the pinwheel of approximately two segments. F. Fidelity scores of BerlinK females (n = 24 flies), BerlinK males (n = 16 flies), empty-Gal4 female (n = 21 flies) executing continuous guided walking between points A and B. G. Number of successes per fly for each genotype with an overlayed barplot. Success is defined as in D. H. Sorted in ascending order by maximum number of *consecutive* successful runs to goal for each fly. Success is defined as in D.

### Olfactory-based remote control of the fly’s walking path

Fruit flies are able to turn towards an antenna that is more strongly stimulated by odors, a behavior called osmotropotaxis (*30*). They will also turn towards the more stimulated antenna when it is triggered optogenetically with light (*34*), with particularly robust movements towards the odor occurring when most (*orco* gene expressing) primary Olfactory Receptor Neurons (ORNs) are stimulated (*35*). Previous studies of natural or fictive osmotropotaxis have experimented with flies that are head fixed on a spherical treadmill; here we extend these studies to remote controlling the walking path of unrestrained flies.

To achieve this, we simultaneously expressed a red-shifted channelrhodopsin, *CsCrimson* (*36*), and a blue-shifted channelrhodopsin, *ChR2* (*37*), under the control of the *orco* gene. (Figure 2Ai). In this case, both red and blue light stimulation will activate *orco* gene-expressing ORNs in both antennae. To create a light-driven, asymmetrical activation of antennas and there by trigger a turn – we painted the right antenna with a pigment that was chosen to pass through red light but not blue, and we painted the left antenna with a pigment that is chosen to pass through blue light and not red (Figure 2B, Supplementary Figure 2). This way red light activates mostly the right antenna, triggering a right turn and blue light activates mostly the left antenna, triggering a left turn. We wrote custom software to track flies walking in an experimental arena, while we automatically trigger turning either with the red or blue light (Figure 2Aii).

**Figure 2:**
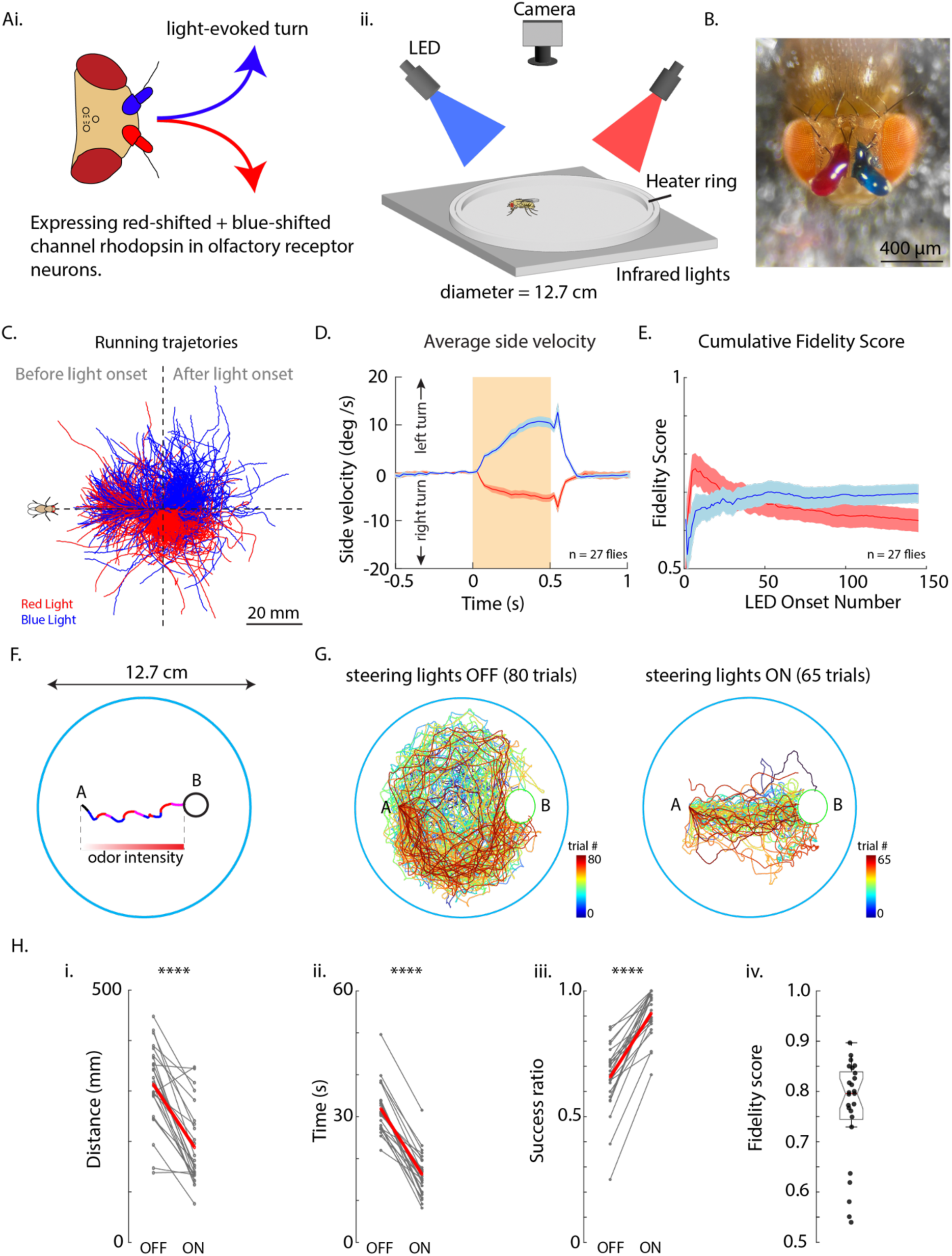
Olfactory based directed turning during free walking. A. Cartoon depicting our experimental setup. (i) We express a red-shifted channelrhodopsin, UAS-CsChrimson and a blue-shifted channelrhodopsin, UAS-ChR2 in olfactory receptor neurons (ORNs) driven by the *Orco*-Gal4 line. We use red (625nm) light to trigger right turns and blue (455nm) light to trigger left turns. Lateralized activation of the olfactory system is achieved by painting antennas with pigments that allow more transmittance of red light on the right antenna and blue light on the left antenna. (ii) Behavioral arena was illuminated with infrared light from below. Tracking camera and stimulation red/blue high powered LEDs were mounted above the arena. B. An example fly depicting antennas painted with red and blue pigments. C. Running trajectories from an example fly, color coded by LED (blue and red), rotated and aligned to LED onset at the center of cross-hairs. To the left of the plot are trajectory segments before LED onset and to the right of the plot are trajectory segments after LED onsets. Fly cartoon on the left shows the flies’ orientation with respect to trajectories. D. Average side velocities and SEM for blue and red stimulated turns are shown (n = 27 flies). E. Cumulative fidelity scores (number of correct turns/total number of stimulations) of red and blue triggered turns as a function of LED onset (n = 27 flies). F. An experimental setup where a fly is guided from a point A in space to an arbitrary goal region B. Real data is shown as an example. Blue segments mark blue LED activation, red segments mark red light activation and purple segments mark when both blue and red lights were on. LED light intensity was scaled with the distance of the fly to the goal. G. (left) Trials from an example fly where LEDs were turned off (80 trials). (right) Trials from the same example fly where LEDs were turned on (65 trials). In the experiment goal region B was randomly selected in space, therefore trials were rotated and aligned at point A for visualization purposes. For a breakdown of individual trials see Supplementary Figure 1. H. Metrics comparing (i) distance traveled (p= 9.68×10^-7^), (ii) time per trial (p=2.02×10^-13^), (iii) success ratio = (number of trials successfully reaching the goal target) / (total number of trials) (p=3.96×10^-10^). Statistics were performed using a two-sample t-Test (n=25 flies). (iv) Fidelity scores for each fly show an average for blue, red and blue+red triggered behavior.

We first tested our ability to drive turning by randomly illuminating flies walking in the arena with either blue or red light. This resulted in walking trajectories of a fly performing turns towards the expected side upon light stimulation (Figure 2C,D). In our analysis of the fly’s proficiency in fictive odor turning, we quantify a ’fidelity score’ – the proportion of accurate turns relative to the total stimulations. Our observations reveal a stable pattern of performance in olfactory navigation across successive stimulations (Figure 2E).

We then examined our ability to guide a fly from an arbitrary point A to an arbitrary target area B in the arena (Figure 2F). With the stimulation lights off, flies essentially display circling or aimless walking behavior, walking to a target area B sometimes by chance. In contrast, when the guiding lights are on, flies show directed walking towards the target area B (Figure 2G). This is demonstrated by comparing ’lights off’ and ’lights on’ conditions, utilizing the metrics of distance traveled, time to goal, and the success ratio of trials – defined as the percentage of instances where the fly reached its goal within 60 seconds (Figure 2Hi-iii). Collectively, our data indicate that we have achieved the capability to remotely control the direction of a fly’s movement, successfully guiding it to a designated spatial point with the flies executing the correct turn approximately 80% of the time (Figure 2Hiv).

### Activation screen to improve odor-guided turning during free walking

We next investigated whether the efficiency of olfactory-guided turning in flies could be enhanced by simultaneous activation of neurons associated with positive olfactory valence (*38*) and olfactory approach behavior (*39*). While flies were guided to walk between two designated locations in space (Figure 3A), we targeted Mushroom Body Output Neurons (MBONs) with red and blue optogenetic stimulation, in addition to activating neurons needed for olfactory guided turning described above (Figure 3B). We employed a fidelity score to gauge the flies’ precision in orienting towards the anticipated direction upon stimulation. Our results showed an improvement in fidelity scores across all tested MBON lines under both red and blue light stimulation. Notably, red light, which penetrates deeper into brain tissue than blue light (*40*), resulted in a significant enhancement of fidelity scores for red light-evoked turns in most MBON lines, including control groups (Supplementary Figure 3A). We interpreted red light as providing ’high’ stimulation and blue light as ’low’ stimulation of MBON neurons. Specific MBON line activation (e.g., MBON-83c) with red light approached the fidelity score performance of visually induced turning (94%), as depicted in Figure 3D, and this high level of performance was maintained throughout most of the experiment (Figure 3E). Heat maps comparing the experimental flies’ movement in control versus 83c conditions indicated an increased occupancy of walking between regions A and B for the 83c group. This study serves as a proof of concept that olfactory guidance in flies can be substantially refined by manipulating specific neural circuits within their olfactory pathway, leveraging both our understanding of the fruit fly’s olfactory system and the precise genetic tools available to target these neurons.

**Figure 3:**
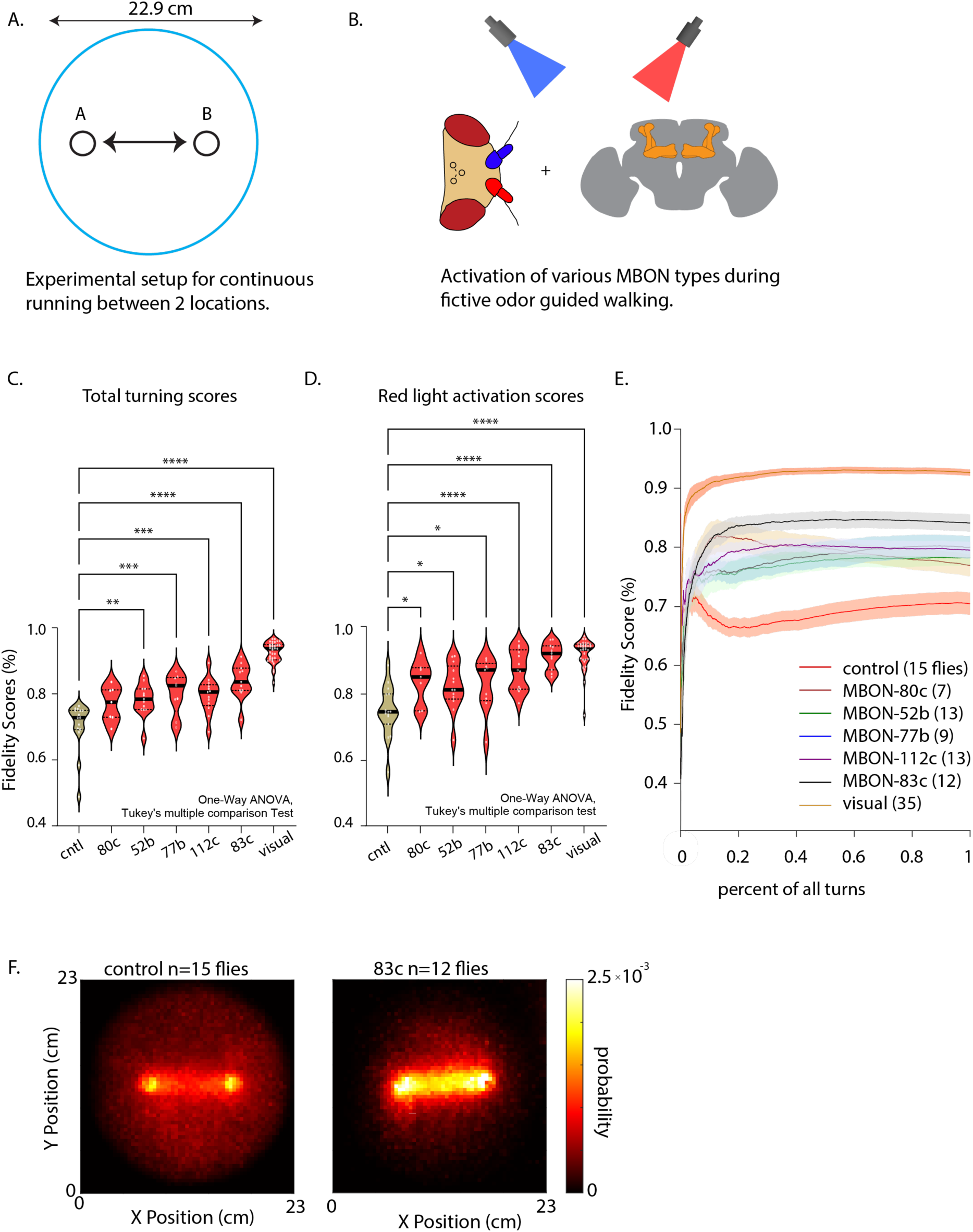
Activation screen to improve olfactory based directed turning during free walking. A. In a circular arena, flies were guided to run between points A and B. B. Our experimental setup was similar to the one depicted in Figure 2, except in addition to asymmetrical olfactory receptor neurons activation, UAS-CsChrimson and UAS-ChR2 were also expressed in 5 Mushroom Body Output Neuron (MBON) types, with one type per experiment. C. Avg left-blue/right-red fidelity scores for each MBON activation condition. Number of * (stars) correspond to significance, * < 0.1, ** < 0.01, *** < 0.001, **** < 0.0001. Statistics were computed using one-way ANOVA and Tukey multiple comparison correction. D. Fidelity scores of red light evoked turns. E. Fidelity scores for each condition as a function of percent of turning (effectively, as a function of time). Number of flies in each condition is shown in parentheses. F. Probability heatmap of fly occupancy in the arena for (left) all control flies (n = 15 flies) and (right) all MBON-83c activating flies (n = 12 flies). All running trajectory data was added together to form a heatmap for each condition.

### Sequencing multiple goals to create complex pattern in ‘writing text’

Building on the visual guidance capabilities of fruit flies, we explored their navigation through a sequence of preset spatial targets, a task analogous to potential engineering applications requiring flies to sequentially visit designated locations. To accomplish this, we programmed a sequence of targets outlining English letters to spell ’HELLO WORLD,’ as shown in Figure 4A. After successfully completing the sequence, the flies were prompted to restart the sequence anew. The typical time to complete the sequence was 17 minutes per trial (Supplementary Figure 6B). Figure 4B illustrates the path of an example fly using our visual guidance system, which enabled all tested flies to precisely ’write’ the phrase ’Hello World’ (Figure 4C, Supplementary Figure 6). For comparison, we present similar patterns executed by olfactory-guided flies, with individual flies ’writing’ each letter (Figure 4D).

**Figure 4:**
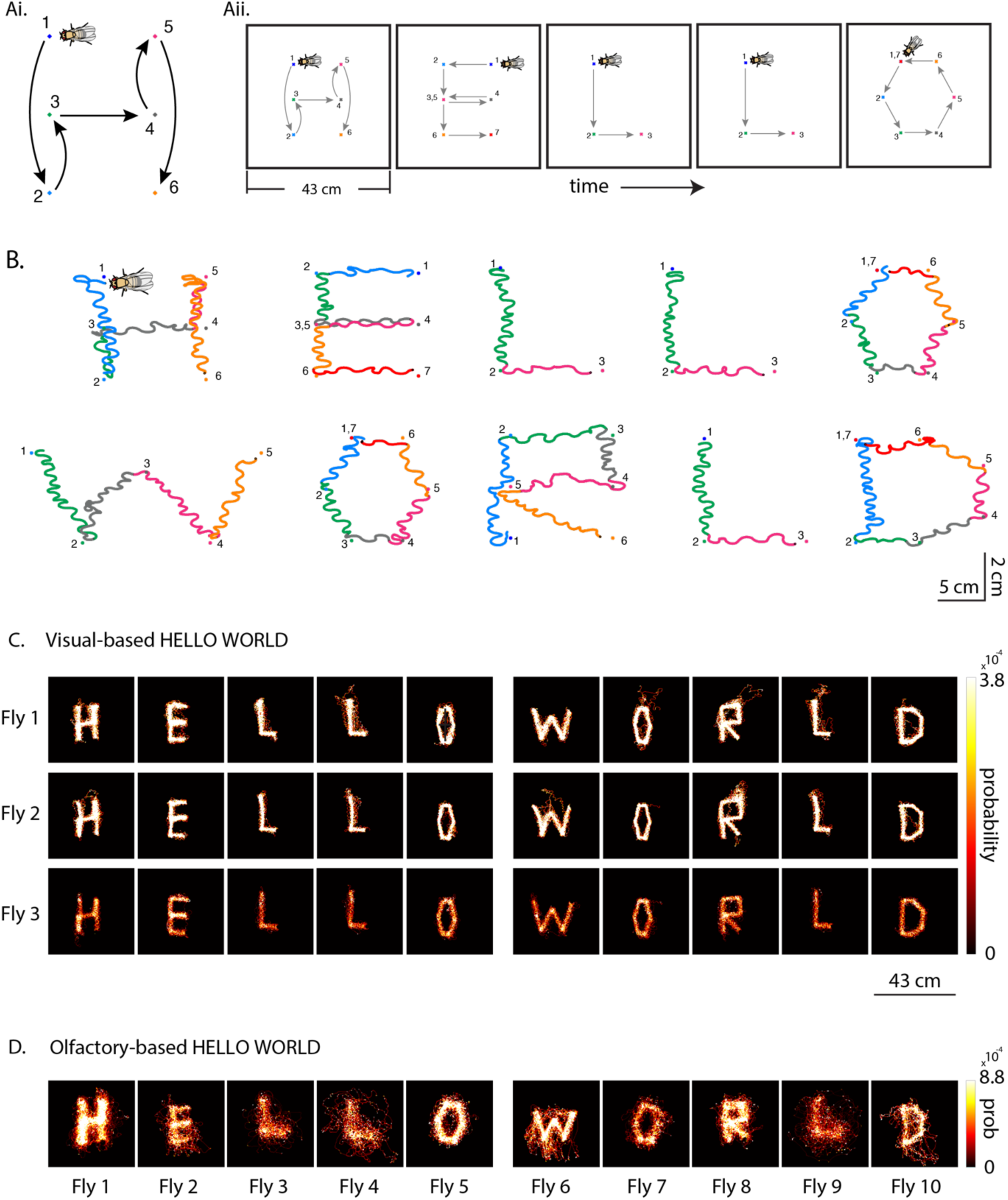
Sequencing multiple goals to create complex patterns in ‘writing text’. A. (i) Schematic depicting the idea of sequencing spatial goals in a pattern of letter H. Numbers and colors represent target goals. Flies’ orientation with respect to the walking trajectory is depicted with a fly cartoon in the top left. (ii) Schematic showing sequencing of patterns to ‘write text’ with fly walking trajectories. B. Example ‘HELLO WORLD’ fly walking patterns ‘written’ by a single fly. Fly cartoon (not to size) shows the flies’ orientation when walking. Running trajectory colors represent runs to a target point (same color as trajectory). Numbers represent target ordinal in sequence. For more examples see Supplementary Figure 4C and Supplementary Figure 7. C. Spatial occupancy probability heatmaps for 3 example flies, showing all running data for each fly. This shows the degree of consistency of ‘writing text’ across letters and across flies. For more examples see Supplementary Figure 6A. D. Olfactory-based “HELLO WORLD” letters. Each letter represents a running trajectory from a different fly. For letters: E, L_2,_ W, D the length of the side of the black square is 43cm. For letters: H, L_1_, O_1_, O_2_, R, L_3_ the length of the side of the black square is 18 cm.

A key benefit of our projector-based system is its capacity for parallel visual guidance of multiple flies, as illustrated in Figure 5Ai. We designed an experiment where each fly functioned analogously to a ‘writing pen’, with the assigned task to independently inscribe the words ’HELLO WORLD’. During the experiment, the assignment of flies to their respective ’pen’ roles could be interchanged, with the constraint that a single ’pen’ represented one fly at any given moment. This approach obviated the need for tracking individual identities of the flies, which is a considerable challenge for real-time feedback experiments. Figure 5Aii depicts three such ’pens’ – each consisting of a unique fly – simultaneously generating three instances of the ’HELLO WORLD’ sequence. While fidelity scores and the precision of guidance were modestly reduced in these multi-fly setups compared to individual-fly experiments, as shown in Figure 5Aiii, the collective performance remained robust, evidenced by the clear replication of the ’HELLO WORLD’ patterns by the group. These findings highlight the potential of our method for swarm micro-robotics applications.

**Figure 5:**
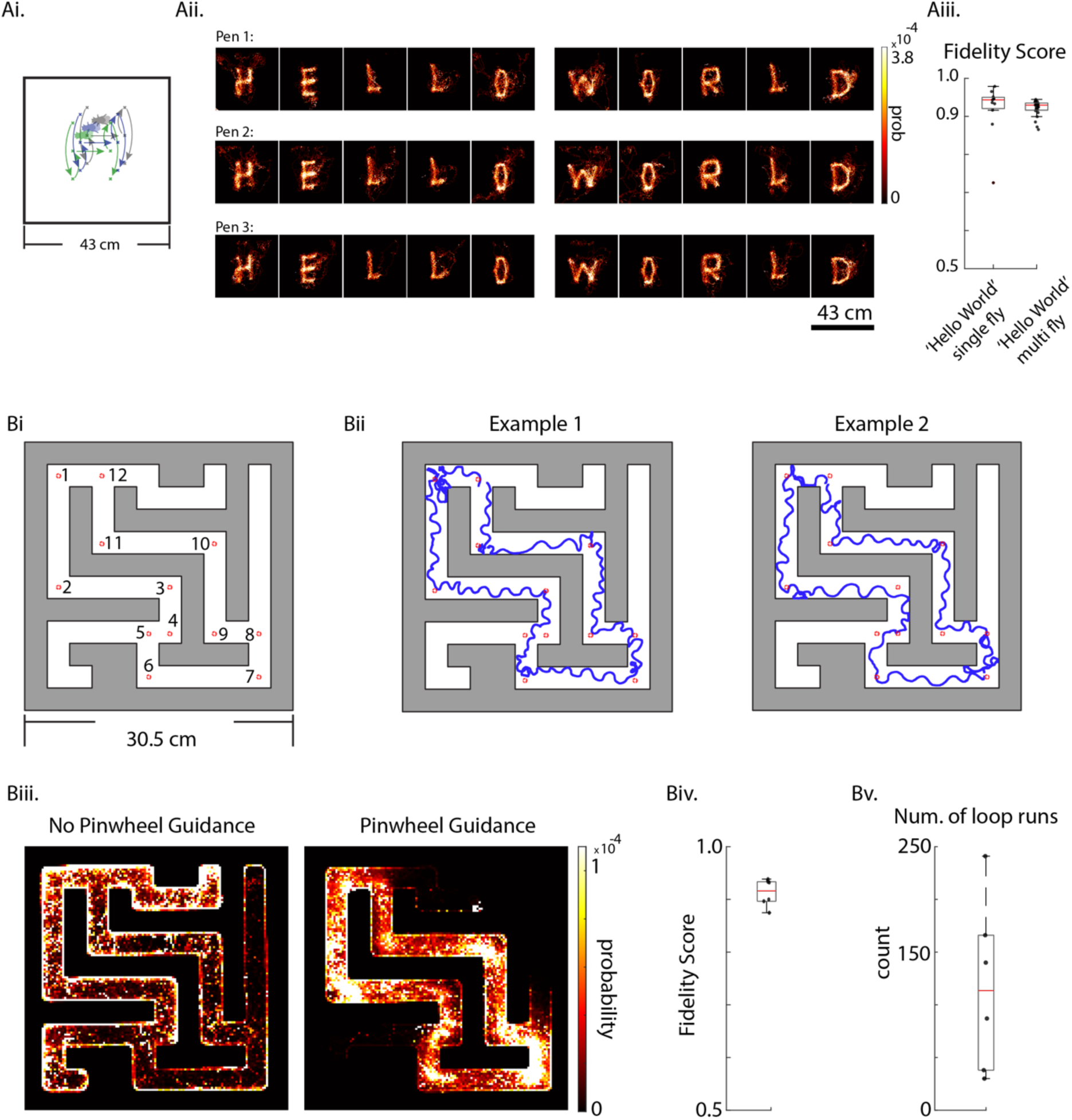
Applications of visually guided turning in freely walking flies. A. (i) Schematic showing multiple flies writing ‘HELLO WORLD’ concurrently, in approximately the same space. (ii) Examples of 3 flies writing ‘HELLO WORLD’ at the same time. (iii) Fidelity scores comparing single fly vs. multi-fly ‘HELLO WORLD’ (p-value = 0.942. n = 11 flies in single experiments, 24 flies in multi experiments (8 experiments, 3 flies per experiment), two sample t-Test). B. Flies visually guided in an engineered environment of a maze. (i) Maze with goal targets in the inner loop. (ii) Two example fly trajectories through the maze. (iii) Spatial occupancy probability heatmaps comparing (left) no pinwheel guidance (n = 4 flies) vs. (right) pinwheel guidance in the maze (n = 6 flies). (iv) Fidelity scores of flies running in a maze for each fly (n = 6 flies). (v) Number of loops that flies ran in a maze, running in sequence goal-to-goal between the 12 goals shown in (i). For examples of individual flies running in a maze, see Supplementary Figure 5A.

### Visual guidance in compartmentalized environments

Building upon the visually guided behavior observed in an open arena, we explored whether flies could adhere to visual cues within the structured confines of a maze – an environment that simulates compartmentalized industrial spaces where flies might be directed to perform tasks. Our maze design incorporated a central ’main route’ with a looping path and multiple dead-end offshoots. Virtual goals were established along the loop, with flies being sequentially navigated through these checkpoints (Figure 5Bi). The flies exhibited a level of navigational precision comparable to that observed in open arena settings, as indicated in Figure 5Bii. Deviations from the main path, such as entering dead-end offshoots, were infrequent, in stark contrast to control experiments without pinwheel guidance, where flies explored the entirety of the maze (Figure 5Biii). The fidelity scores for maze-based guidance were comparable to those from open field experiments (Figure 5Biv). Flies consistently followed the main route for hundreds of iterations (Figure 5Biv). These findings collectively affirm that the visual guidance of flies is equally effective in restricted environments.

### Flies guided to carry a load between two spatial locations

Exploring the potential for fruit flies in robotic applications, we considered their capacity for transporting small cargo across designated spatial locations. We asked, what is the maximum weight that a fly can carry while still effectively navigating between two spatial locations under visual guidance? To investigate this, we affixed incremental weights to the thoraces of fruit flies and directed them between two points, analogous to the procedure depicted in Figure 1A. Fidelity scores (Figure 6C) and spatial heat maps of their trajectories (Figure 6B) show that the flies reliably transported loads up to 1.1 milligrams, similar to the weight of the fly (Supplementary Figure 5B). Performance declined with additional weight (Figure 6C-E); however, some individuals were capable of repeated linear navigation even with the maximum weights (Figure 6D). Remarkably, certain flies managed to carry loads of 0.9 milligrams or less across distances reaching 500 meters, equivalent to approximately 200,000 times their body length (Figure 6D).

**Figure 6.**
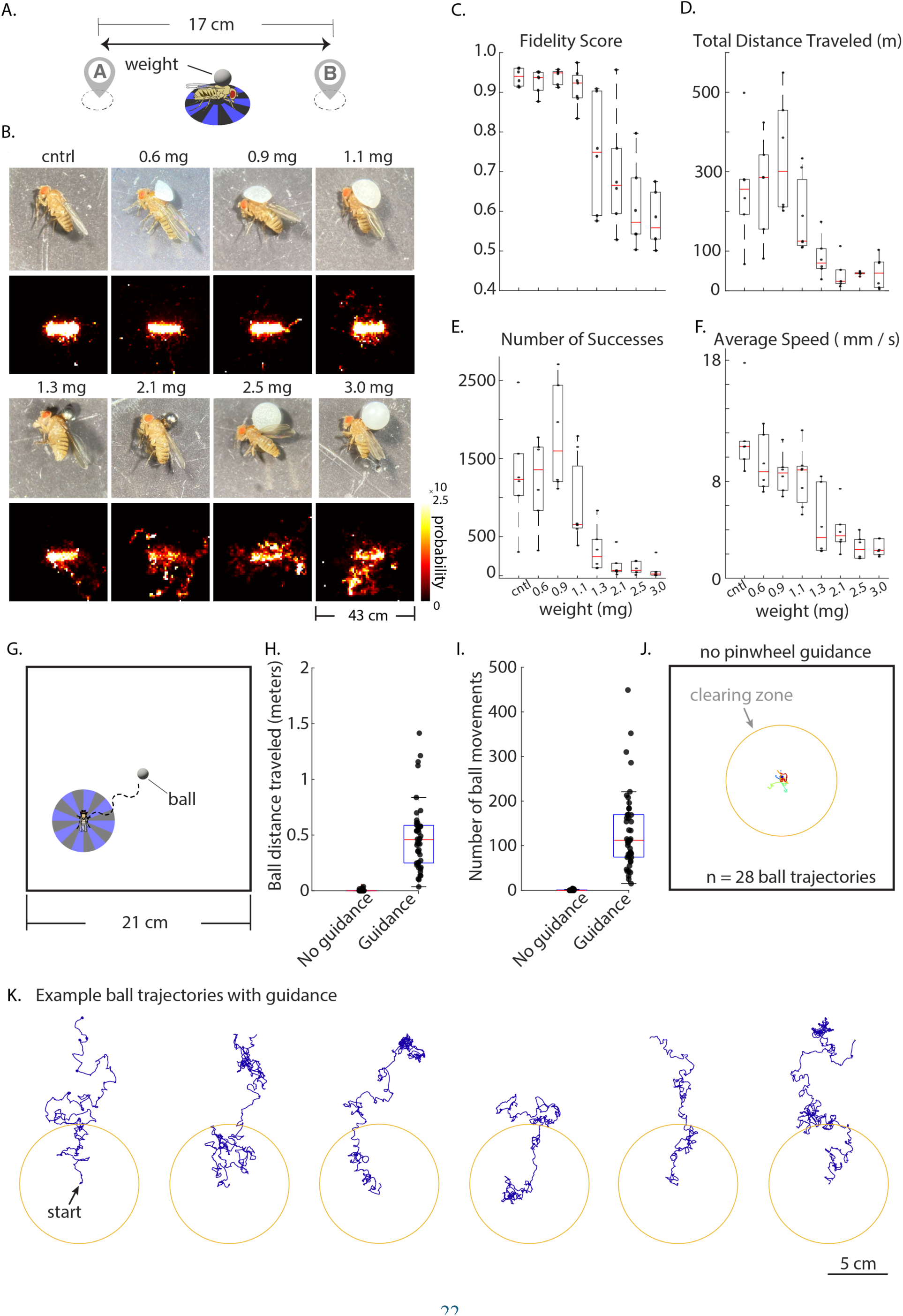
Example micro-robotic applications: 1) Flies moving weight across two designated locations and 2) Flies relocating a spherical object away from a designated location. A. Schematic of the experiment. Experimental design is similar to Figure 1A, except flies are carrying weight of varying magnitude attached to their thorax. B. (Top row) Images of flies with weights of different magnitude glued to the thorax. (bottom row) Spatial occupancy probability heatmaps for each weight condition. (control: n = 6flies; 0.6mg: n= 6 flies; 0.6mg: n= 6flies; 0.9mg: n= 6 flies; 1.1mg: n= 7flies; 1.3mg: n= 6 flies; 2.1mg: n= 6flies; 2.5mg: n= 6 flies; 3.0mg: n= 6 flies) C. Fidelity scores for each weight condition presented as boxplots. Note, there is a sharp drop off in performance after 1.1mg. D. Total distance that a fly traveled carrying weight for each condition, presented as boxplots. E. Number of successful trials per fly. Success is defined as reaching a goal within 60 seconds. F. Average speed of flies carrying weight for each condition, presented a boxplots. G. Schematic of the experiment, showing a fly visually guided to a small spherical object (10 mg ball). H. Total ball distance traveled for each experiment, comparing conditions without (n=28ball trajectories) and with (n=47 ball trajectories) visual guidance (p-value: 8.07×10^-7^). I. Number of ball movement events for each experiment, comparing conditions without (n=28ball trajectories) and with (n=47 ball trajectories) visual guidance (p-value: 3.31×10^-9^). J. Ball trajectories when no visual guidance is provided. Note, there is minimal ball movement (n = 28 ball trajectories). The circle depicts a clearing zone of 5 cm radius. K. Examples of partial ball movement trajectories caused by flies repeatedly visually guided towards the ball. Ball trajectories were truncated for clarity.

### Flies guided to relocate a spherical object: implications for robotic clearing applications

To assess the potential of biological agents to augment robotic cleaning operations, we explored the ability of flies to transport a spherical object significantly larger than themselves. This inquiry is pertinent to the development of robotic systems engineered to autonomously clear debris from designated areas. We manipulated the flies’ behavior by directing them toward the spherical object (10 mg) and deactivated visual cues when a fly approached within 5 mm of the object (Figure 6G). The flies interacted with the object typically 110 times (Figure 6I), achieving a median displacement of 49 cm (Figure 6H). Our results indicate that visual cues are crucial for the flies’ effective interaction with the object; without these cues, engagement was minimal (Figures 6H-J). Trajectories of the flies that moved the object at least 5 cm beyond the center of the arena are illustrated (Figure 6K), with 85% of flies able to perform such clearing (n = 47 flies). This finding not only underscores the flies’ capacity to manipulate object placement but also suggests a scalable strategy for integrating biological elements into robotic systems to enhance operational efficiency.

## Discussion

In this study we demonstrate the ability to remotely control the heading of a walking fruit fly using two separate methods that are based on visual or olfactory guidance. Each of these methods has unique advantages and disadvantages. The olfactory-based approach benefits from a straightforward stimulation process. To our knowledge, this study is the first reported instance of blue and red light activation of behavior in the same fly (*11*). Conversely, the visual stimulation method circumvents genetic and physical manipulation, providing a non-invasive alternative.

Nevertheless, it demands a sophisticated setup utilizing projector technology to effectively deliver visual cues. Previous studies have demonstrated the manipulation of behavior of freely walking or flying flies in virtual environments, highlighting the potential of this approach (*41, 42*). We also demonstrate that flies can carry weight for considerable distances in relation to their body length and follow guidance cues to walk through a sequence of spatial goals in a constrained environment. This opens doors to engineering applications where a small weight (e.g. something that weighs 1 mg) needs to be transported from specific engineered compartments.

In recent years, the integration of biological organisms with robotic systems has gained considerable interest in the field of bio-robotics. Several studies have demonstrated the utility of small insects in robotic applications, exploiting their innate behaviors and physiological features to enhance robotic functionalities. Notably, the cockroach, *Gromphadorhina portentosa*, was utilized to develop bio-hybrid robots controlled via electrical stimulation, showcasing the potential for insect-driven locomotion in robots (*43*). Similarly, beetles, *Cotinis texana*, were explored for micro-robotic applications, implementing a miniature wireless control system to direct their flight paths (*44*). These pioneering efforts have set a precedent for our current work with a considerably smaller insect, *Drosophila melanogaster*, where we propose a novel micro-robotic platform that harnesses the unique genetic tools and neurobiological knowledge of fruit flies to advance the field of micro-robotics.

In addition, advances in optical microcircuit technologies (*45*) open the possibility for control of flies outside the laboratory environment in a biohybrid design (*46*). This can be accomplished via engineered ‘backpacks’ that will be carried by flies. These backpacks can potentially provide light stimulation, both in the context of olfaction via antennae stimulation or vision via moving visual patterns. Such an autonomous biological robot would be the basic unit in a swarm and potentially able to address the long-term visions in micro-robotics including search and rescue, pollination or environmental monitoring applications.

In addition, usage of this technology extends beyond engineering utility, offering insights into the neural circuitry of fruit flies. Our examination of mushroom body output neurons (MBONs) reveals a marked increase in directional accuracy upon activation, particularly evident in the GABAergic subsets, MBON-γ3, γ3β’1 (MBON-83c), and MBON-γ1pedc>α/β (MBON-112c). Notably, the activation of GABAergic MBONs sustains precision in navigational control over extended periods. We hypothesize that the pronounced effect of MBON-γ3, γ3β’1 may be linked to their role in consolidating ’safety memory,’ influencing the selection of ’safer’ behavioral choices (*47*). This suggests that associative mechanisms might underpin the persistent selection of the induced odor as a navigational cue.

Flies persisted robustly in following our guidance cues for many hours despite the absence of explicit rewards in our task reinforcing this behavior. This encourages both engineering applications and raises scientific inquiries that could uncover the neural mechanisms behind such persistence. The technology described in this study opens the potential for a general set of experiments where guidance cues can conflict with environmental features, thus uncovering latent behavioral preferences of the fly.

Our era boasts an array of genetic tools for neural intervention and a complete brain circuitry atlas. Despite these advancements, our capacity to delve into the fruit fly’s natural behaviors and decision-making processes has mostly been limited to passive observation. The ability to precisely position flies in space helps overcome these constraints, empowering researchers to initiate novel studies that explore how fruit flies navigate, perform social interactions, and interact with their environment. This enriches our understanding of their innate behavioral responses while also leveraging these insights in future micro-robotic applications.

## Acknowledgments

We thank Rosy Hosking, Andrew Murray, Pallav Kosuri for helpful comments on the manuscript. We thank Yoshi Aso, Avalon Owens, Kevin Gozzi, Buck Trible, Paul Shamble, Leigh Needleman, Winfield Hill, Michael Burns, Andrew Murray for helpful discussions. We thank Erik Madsen for machine shop support. We thank Scott Bevis and John McKenna for help with instrumentation prototyping.

## Author contributions

Conceptualization: CN, CS and AR

Methodology: KI, CN, CS, AR

Investigation: KI, CN, AR

Visualization: KI, CN, AR

Funding acquisition: AR

Project administration: AR

Supervision: AR

Writing – original draft: KI, CN, AR

Writing – review & editing: KI, CN, CS, AR

## Data and materials availability

All data and code will be available in a public database.

## Materials and Methods

### Fly strains & Maintenance

Flies were kept in a vial of standard cornmeal fly food (Archon Scientific, Corn syrup-soy medium) with 12: 12 hr. light: dark cycle inside a 25 °C incubator (Insect Environmental Chamber, Biocold Environmental Inc.) with humidity set to 55%. Flies that were tested optogenetically were raised with 2.3 mM all trans-Retinal (Sigma-Aldrich, Cat. No. R2500) in vials wrapped with an aluminum foil sheet to avoid light exposure. Adult flies were aged based on the number of days since the time of eclosion. Flies with *Orco-Gal4 > UAS-CsChr, UAS-ChR2/empty-Gal4* were used for olfactory optogenetic experiments for both short line (point A to B walking) experiments as well as letter spelling experiments. For the MBON screen experiments, virgin flies with *Orco-Gal4 > UAS-CsChr, UAS-ChR2* were crossed with mushroom output neuron Gal4 drivers (Table S2) that target output neuron groups known to alter flies’ response to olfactory stimulation (*38, 39, 48*) and empty-Gal4 (Stock# 68384, BDSC) for a control. Berlin-K (Stock# 8522, BDSC) and *empty-Gal4* adult flies were used for visual guidance experiments. Genotypes of adult flies used for each figure in the manuscript are described in Table S1 below.

### Fly housing, breeding, and selection for experiments

Once a fly was selected for an olfactory experiment, it was painted and then moved into a separate vial by itself, where 200 µl of the 35 mM ATR stock solution was poured into kimwipes wetted with 3 ml water. The fly was then kept in the dark (to not degrade the ATR) and without food for 12-24 hours until it was used for an experiment. For visual experiments the fly was selected and added to the arena for the experiment without being painted or starved first.

### Setup of a larger arena for Figure 1

In Figure 1 we used a large square arena, 43cm on each side, with an infrared light source underneath the arena (Knema, 18 x 18-inches edge lit panels, IR 850nm LEDs, knema.com, South Korea). Along the edges ran a heating element (McMaster, 2156T71) to discourage flies from resting on the edges, and remain mobile in the arena. A FLIR camera (Blackfly S BFS-U3-63S4M) was positioned above the arena to track the position of the fly. Along with a projector (Optoma ZH406 (deprecated) or Optoma ZH450) to project pinwheel stimulation patterns onto the flies’ walking surface.

### Calibration of projector for Figure 1

In our system, we track the position of the fly in camera coordinates (as an XY pixel position in the image). To project an image centered on the fly, we needed to generate a mapping between camera coordinates and projector coordinates. This was done by projecting an array of crosses with known coordinates in ‘projector space’ whose centers were easily discernible by the camera. We then take their camera coordinates and map the known coordinate in ‘projector space’ to ‘camera space’. This array of crosses was shifted around to tile the whole arena, and then a smoothing algorithm was run to create an entirely smooth mapping of points. This method allowed us to have a direct camera-to-projector measurement for each point in the physical space of the arena and resulted in an accurate placement of the pinwheel, centered at the position of the fly in the arena.

### Pinwheel parameter selection

We detected the fly using a similar thresholding method as described above. The pinwheel was centered on the position of the fly and consisted of alternating blue and black sectors. We rotated the pinwheel either clockwise or counterclockwise around the fly to guide the fly to turn either right or left respectively. For spatial frequency we chose 16 segments (22.5° per wedge) and our rotational velocity ranged from 240-360°/second – fitting well into a range defined by previous studies (*32, 41, 49*).

### Software for running visual guidance experiments in Figure 1

In the following set of experiments a sedated fly was placed in the center of the arena to initiate the experimental procedure. The objective was to navigate the fly between two points, set 13 cm apart, within the arena. Success was defined as the fly reaching a goal region, delineated as an area within 1.9 cm of the targeted point. Upon achieving this, the trial was deemed successful and a subsequent trial commenced, with the destination being the alternate goal point. Trials were limited to a 60-second timeframe. In instances where the fly remained immobile – failing to move beyond 1.25 millimeters within a 0.25 second period – for 15 seconds, or if the goal was not reached within the allocated time, the trial was concluded and marked as unsuccessful.

Control over the flies’ navigation was exerted through a pinwheel mechanism, the rotational speed of which was modulated in response to the fly’s heading angle relative to the designated goal point. A heading within 60 degrees of the goal point resulted in a rotation speed of 240 degrees per second. As the deviation from the goal point increased to between 60 and 90 degrees, the rotation speed was adjusted to 1.1 times the base speed, equating to 264 degrees per second. For deviations between 90 and 120 degrees, the speed was increased to 1.2 times the base speed, or 288 degrees per second. For the most substantial deviations, ranging from 120 to 180 degrees from the goal, the rotation speed was increased to 1.5 times the base speed, achieving 360 degrees per second. This graduated scaling of speed was implemented with the intention of expediting the correction of flies’ larger deviations from the intended path.

### Painting procedure for Figure 2 and dyes used

For olfactory experiments in Figure 2, a 1-3 day old virgin female was placed in a 5.0 ml culture tube (VWR, Cat. No. 60818-565) in an ice bath for approximately 30 seconds until the fly was completely immobilized. The anesthetized fly was then placed into a custom holder made of 0.001" stainless steel sheet (McMaster: 3254K311) with a small hole to expose the anterior head of the fly for painting. The fly’s thorax was glued to the metal holder to immobilize it with light-cured adhesive (Henkel Loctite, Cat. No. AA3972). We used two pigments, both food coloring powders; the blue pigment (LorAnnOils.com, Lorann, Blue Powder 1310-0400 Lot: Z0958), and the red pigment (Jelife, Red Food Color Powder; GuangDong Province, China). These pigments were chosen for their transmittance of specific wavelengths. Several candidate pigments were tested on a spectrophotometer (Agilent Technologies, Cary 5000 UV-Vis-NIR) and these two pigments were selected because there was no overlap in their transmittance. The locations of the peaks in transmittance aligned well with the activation of the blue shifted and red shifted channelrhodopsins (455 nm and 625 nm for blue and red respectively), as seen in Supplementary Figure 2. These pigments were mixed into UV curing glue (Henkel, Cat. No. AA3972) to form a paint, with an approximate mass ratio of 35% of the red pigment to the glue, and 15% for the blue pigment. The right antenna was painted with the red dye mix and the left with blue dye mix. As a ‘paintbrush’ we used micropipettes pulled from 0.8-1.1mm borosilicate glass tubes. The dye mixes were cured by exposing the fly’s head to a UV light (Electro-Lite, LED-200) for a few seconds. After painting the antennae, we carefully peeled off the glue securing it in the holder and collected the fly in a 5.0 ml culture tube (VWR, Cat. No. 60818-565). The tube was then put on ice and immobilized the fly in order to insert into the arena for experimentation.

### Painting of flies for the MBON screen and the olfactory HELLO WORLD assay

This painting procedure followed all of the above, as in Figure 2, with the addition that both eyes of the flies were painted with a black paint mix to make them effectively blind. The black pigment for this paint mix was from Jelife (Black Food Color Powder; GuangDong Province, China) and mixed into the glue (Henkel, Cat. No. AA3972) for a mass ratio of 30%. After the painting, the flies were kept in a plastic vial on 2.3 mM ATR (Sigma-Aldrich, Cat. No. R2500) until the time of testing (200 µl of the 35 mM ATR stock solution was poured into kimwipes wetted with 3 ml water).

### Setup of arena and imaging for olfactory guided experiments

The experiments in Figure 2 were carried out in a circular dish made of Delrin acetal resin sheets (McMaster, 8573K122) and with a diameter of 12.7 cm. Along the edge of the circular rig we ran a heating element (Pelican Wire Company, Resistance Wire, Advance Alloy, 32 AWG) set to 40°C in order to keep the flies from standing along the edge of the arena. The top of the arena was composed of a clear sheet of (McMaster: 8536K144) to ensure the flies would not fly away, and coated in SigmaCote (Sigma Aldrich, SL2-25ML) to prevent the flies from being able to walk upside down on the cover. The arena was backlit by an infrared light panel (Knema, LED 850 nm 6×6”, knema.com, South Korea) placed 4 cm below the arena. We used a Blackfly S (FLIR, Blackfly S BFS-U3-13Y3M) camera to track the position of the fly. The camera mounted a lens (ArduCam, 2.8-12mm varifocal c-mount lens LN049) and an infrared longpass filter (Hoya, 46mm infrared R72 filter). Two LED lights, one red and one blue (ThorLabs, M455L4 and M625L4), emitting narrowly at 425 nm and 625 nm respectively to minimize any crossover in their emission and to avoid unintended activation of the wrong channelrhodopsin, were positioned such that there would be even illumination for both red and blue light across the entire arena.

### Details of fly tracking and stimulation algorithms

The experiments illustrated in Figure 2 consisted of tracking the fly using a background subtraction method and thresholding of the resultant image to determine the exact pixel location of the fly. Before the start of each experiment, a reference background image was saved and subtracted from each new image. The resultant array was then thresholded, leaving only the points where the fly was currently located.

For parts C, D, and E the experiment consisted of trials where any time the fly was walking with a minimum velocity of 1.0 cm / second and it had been at least 8 seconds since the previous trial, either the red or blue light (randomly chosen) were turned on for 0.5 seconds and the trajectory of the fly was recorded. This was to ascertain the fidelity of turning exhibited when the fly was illuminated with either red or blue light, and to examine the behavioral response to each color of light enabled steering control of the fly’s walking behavior.

For parts F, G, and H trials were initiated any time the fly was near the edge of the arena and started to move away from the edge (so that it would have enough space to run forwards), as long as there had been more than 30 seconds since the last trial. At the initiation of a trial, a goal region was designated 6.82 cm across from the fly’s current position. If a fly came within 1.9 cm of the goal region center, the trial was considered successful. The fly’s current heading was calculated each frame by subtracting its current position from its position 5 frames beforehand. If the current heading was pointing within the goal region then both red and blue lights were turned on. If the heading of the fly lay to the left of the goal, the red light as automatically turned on to attempt to correct the heading by triggering a right turn. If the heading pointed to the right of the goal the blue light was turned on to trigger a left turn. The light intensity was scaled with the distance of the fly from the goal. The lights would get brighter the closer the fly got to the goal, with blue light from 0.5 mW to 2.5 mW, red light from about 0.3 mW to 1.67 mW. Lights were turned off if the fly stopped moving for 1 second and if the fly didn’t move for 15 seconds the trial would terminate as a failure. Each trial ran for 60 seconds or until the fly reached the goal, whichever came first. If the fly had not reached the goal within 60 seconds, the trial was terminated and classified as a failure. A new trial was initiated after the intertrial period of 30 seconds. Information regarding the fly’s current position, the state of the lights, and the state of the trial was saved and exported to a .mat file every 30 seconds for later analysis. Control trials were randomly interspersed with experimental trials, where the position of the fly was recorded and a goal point was set, but the lights were off, to record the fly’s natural behavior when not being guided by lights.

### Explanation of fidelity scores

The fidelity score metric that is displayed in Figure 1F, 2 E and H, as well as later in the paper, is a simple metric that demonstrates how faithfully the fly was turning in response to the stimulus. It is a fraction of the number of successful turns in the direction indicated by the stimulus, divided by the number of times we provided the stimulus to attempt to guide the fly in that direction. In other words, it is a sum of the number of times the fly turned left when the blue light was turned on, plus the number of times the fly turned right when the red light was turned on, divided by the total number of instances we provided stimuli to the fly. In the case of visual guided experiments, stimuli in this equation are from clockwise and counterclockwise rotations of the pinwheel. We measure each potential turn by computing the fly’s side velocity during a specific stimulation period. Once the trajectory was rotated to align it with the x-axis in the positive direction, left turns occurred if the fly’s average side velocity was >0 during the stimulation period, and <0 for right turns.

To calculate the cumulative fidelity score as seen in Figure 2E, the fidelity score was computed at each turn throughout an experiment by dividing the number of successful turns by the total number of times the stimulus was triggered. This plot includes the mean fidelity score, as well as the standard error of the mean in the lighter shaded region around the mean.

### Experimental setup of a larger circular arena for Figure 3 (MBON screen)

In Figure 3, the circular arena setup was expanded to be a larger circle with a diameter of 20cm, and with additional infrared lights (SMD3528-300-IR Infrared (850nm) LED Light Strip, part number: 3528IR850NM-60B/M-5M) added underneath to ensure even illumination of the wider arena. Two of each ThorLabs LED were now used to increase illumination over the larger area, scaling from about 1.02 mW to 5.12 mW for blue, and 0.55mW to 3.33mW for red. Along the edge of the circular arena we ran a heating element cable (McMaster: 2156T71) to ensure that the flies would stay moving in the middle of the arena. The two goal points were 4.5 cm apart in fixed locations and centered on the middle of the arena.

### MBON activation screen to improve olfactory guidance

Activation of different groups of mushroom body output neurons (MBONs) affects how flies respond behaviorally to sensory stimulation (*38, 39*). To determine whether MBONs can modulate odor-mediated navigational control, MBONs that elicit a significant positive valence response upon stimulation, MB052B, MB077B, MB112C, MB083C (*38*) and the MBON that modulates how flies persist in responding to olfactory stimulation, MB080C (*39*) were selected (Table S2) to conduct a mini screen to identify MBONs that enhance performance of odor-mediated navigational control. To identify a MBON driver that enhances the navigational performance, fidelity scores of each MBON group were calculated as above and compared to that of the control flies (flies with Empty-Gal4 crossed to flies with *Orco > CsChrimson, ChR2*). Statistical analysis of the acquired data (One-Way ANOVA, Tukey’s Multiple Comparison Test) was conducted using Prism software.

### Generation of HELLO WORLD letters

Each letter of “HELLO WORLD” was designed in MATLAB, containing multiple goal points which correspond to x, y coordinates such that each letter consists of the smallest number of straight lines needed to make it legible. The order of the goal points was determined to ensure continuous movement to completion. When a fly reached within 4 mm of the correct goal point, the fly was guided to the next ordered goal point. Once the fly completed one letter, it was guided to the first goal point of the next letter. Thus, the fly was navigated through each of the goal points to complete spelling each letter of “HELLO WORLD”.

### Software design for HELLO WORLD experiments

To carry out ‘HELLO WORLD’ experiments, we wrote custom python software that selected arbitrary letter patters and sequenced through spatial goals in order defined by these patterns. After running through all the letter patterns, and upon completion of the last goal point in the last latter, the letter sequencing would restart from the beginning. This software was also used to generate letters in Supplementary Figure 4C.

### Olfactory LED stimulation code running ‘HELLO WORLD’ experiments

Similar to patterns of letters sequencing for projection experiments, we sequenced patterns for olfactory experiments. In this case, the turning signal was driven through red and blue LEDs as described above, instead of the pinwheel patterns driven by the projector.

### Figure 4D Olfactory HELLO WORLD

To demonstrate how well flies can be guided spatially using olfactory cues, flies empty-Gal4 (*UAS-CsChrimson; Orco-Gal4; UAS-ChR2/empty-Gal4)* or MB080C (*UAS-CsChrimson; Orco-Gal4; UAS-ChR2/R50A05-GAL4.DBD, R33E02-p65.AD*), with their antenna painted as described above, were optogenetically guided to spell each letter of “HELLO WORLD” (n = 4-17 flies per letter). The best performance for each letter was selected and combined to make the best “HELLO WORLD” (Figure 4D). In this figure, each letter was ‘written’ by a different fly.

### Setup of multi-fly experiments

The projector-based arena was the same for multi-fly experiments as for single fly experiments. Control code was modified to track multiple flies in the same image, without tracking their individual identities. At every time step, each discovered fly in the image was then assigned to a ‘pen’ – a manager that keeps track of goals in each pattern for each letter in one of the ‘HELLO WORLD’ instances. In these experiments, there were 3 pens that were managing running trajectories of 3 flies. Fly tracking data from each pen was then saved separately for further analysis. The maximum number of flies that could be processed was constrained by the duration of each time step (running at 16 frames / second) required by our prototype code for real-time detection of the flies and production of the turning signals.

### Description of maze design

The maze was designed in MATLAB to contain the “main route” that flies can be visually guided to walk through. Multiple dead ends were added to the maze to evaluate the flies’ ability to follow the visual guidance without distraction. Goal points were added to corners of the main route to enable targeted guidance of the fly through the maze. The actual maze (30 x 30 cm) was made of acrylic (McMaster, 8505K817) with each individual path 2.5 cm wide and the wall 3.5 mm high.

### Weight carrying assays

To determine how much weight a fly can carry on its back while following visual navigational guidance, weights of different masses (0.6 mg, 0.9 mg, 1.1 mg, 1.3 mg, 2.1 mg, 2.5 mg, and 3 mg; Table S3), were mounted and glued, with a thin layer of light cure adhesive (Henkel, Cat. No. AA3972), onto the dorsal thorax of the fly. A layer of the adhesive used per fly weighed less than 100 nanograms. The diameter of the balls ranges from 0.6 mm to 1.6 mm. To generate a range of smaller weights, ball fragments were generated by splitting 2.5 mg Nylon balls (McMaster-Carr) to desired weights. Flies with weights on their backs were guided visually to walk between two goal points that were 17.14 cm apart. When a fly reached within 1.5 cm of a goal point, it is guided to walk to the other goal point.

### Assisted fly-ball interaction assay

The larger behavioral test arenas (45.72 x 45.72 cm) were divided into four quadrants, each of which is separated from adjacent quadrants by 8 mm-plastic dividers, to record interactions between male flies and small white Delrin Acetal Resin Balls (McMaster-Carr: 9614K51). The balls were 2.38 mm in diameter and 10 mg in weight. Some rigs had heat wires around the outer side walls to prevent flies from crawling up the walls. The recording of the interactions was done at 16 frames per second. The guidance pinwheel (2.4-2.6 µW/mm^2^ at 480 nm) was projected by an Optoma projector (Optoma ZH450) centered at the location of a fly in each quadrant. Before each experiment, balls were placed in the middle of each quadrant and flies in inner or outer corners of the quadrants. Male flies were reared in a vial with fly food (Formula 4-24, Caroline Biological Supply Company or Corn Syrup/Soy food, Archon Scientific) for one day after eclosion prior to experiments. Flies were guided repeatedly to walk toward balls by the pinwheel guidance throughout the experiment. The pinwheels were turned off when flies were within a 5 mm radius of the balls. In analysis of ball movement data in Figure 6H-I, ball movements less than 1.5 mm were removed to account for tracking noise when flies are close to the ball.

### Analysis of olfactory guided walking direction (Figure 2)

To look at each individual turn, as in Figure 2 C and D, we took snippets of time just before the lights turned on and for 1 second after the lights were turned on to capture the heading of the fly throughout the potential turn. Rotational and translational transforms were applied to the trajectories to align them such that the trajectory at the onset of the light was centered at (0,0) and the trajectory for 0.25 seconds before the onset was aligned with the x axis. This allowed us to compare turning behavior after the onset of light.

### Analysis of Hello World ‘writing’ data (Figure 4)

Figure 4 B was created by finding one example fly’s trajectory from which to select a continuous trajectory of one iteration of the ‘HELLO WORLD’ experiment. This trajectory was then segmented by pattern (letter), with the paths from the last goal of one letter to the first goal of the next discarded. The segments of the trajectory leading to each of the individual goals that compose a pattern were then illustrated with a different color. The best visually clean letters were chosen for this example fly. Similar procedure was applied to writing trajectories in Supplementary Figure 7, except the letters shown are in continuous order of writing.

Figure 4 C was generated by taking three flies’ entire experiment trajectory, and then segmenting out the data based on the latter pattern. This trajectory data was then turned into a heatmap by using the binned probability that the fly would be in any area over the course of the experiment. Some individual runs have been excluded from this data to enable us to clearly display each letter that the fly was ‘writing’ in order, as a summary of the whole experiment. If the number of failures for a particular instance of a letter exceeded 10 or if a given letter’s trajectory moved by more than 12.5 cm from the line between two goals, then that one instance of a letter was excluded. The fraction of letters accepted for display is shown in Supplementary Figure 6D.

**Fig. S1.**
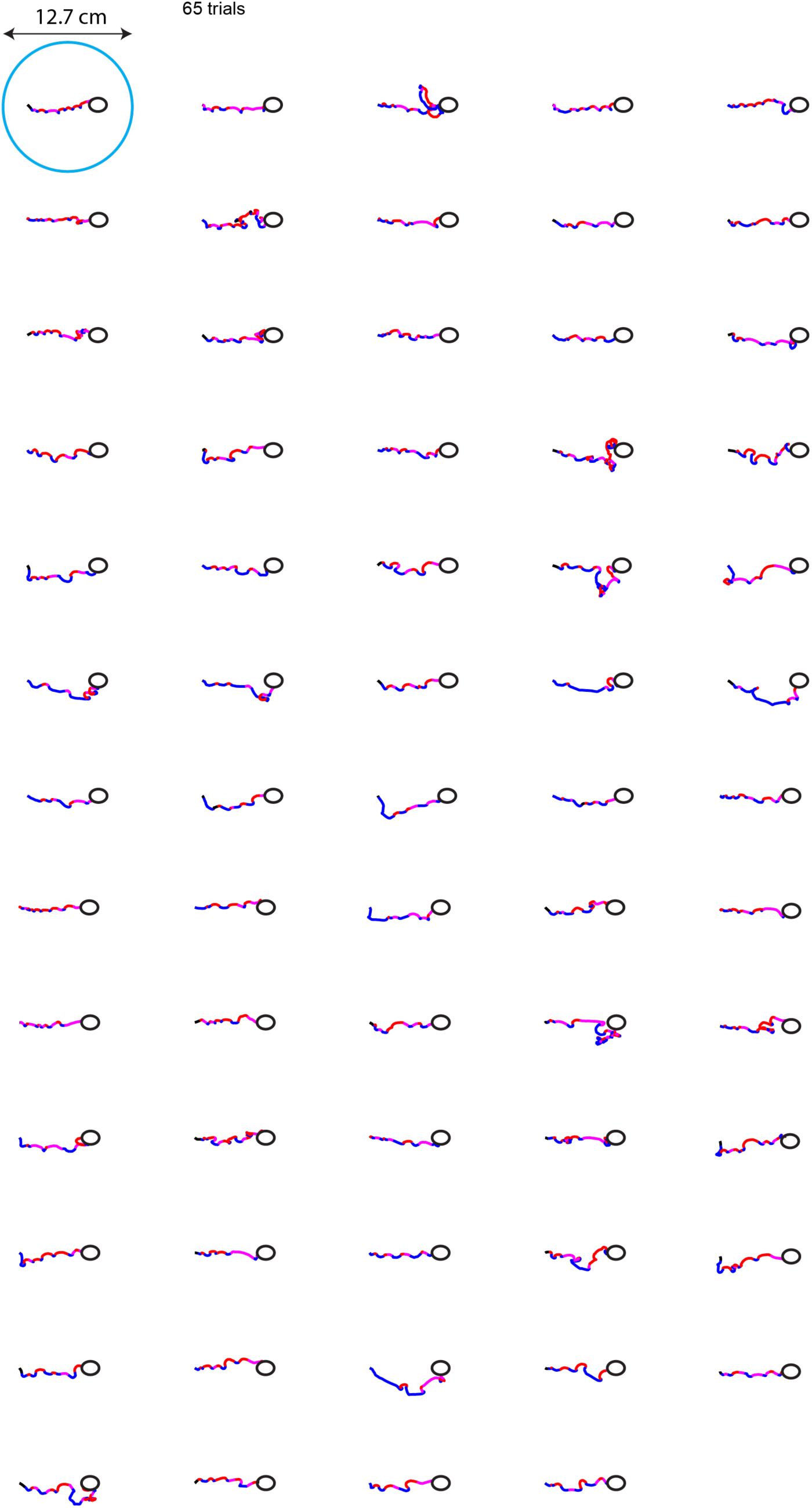
Individual running trajectories from the example fly in Figure 2G. Color scheme is the same as in 2F. Red and blue indicate red and blue light illumination during a run respectively. Purple indicates that both lights were turned on to encourage the fly to run straight.

**Fig. S2.**
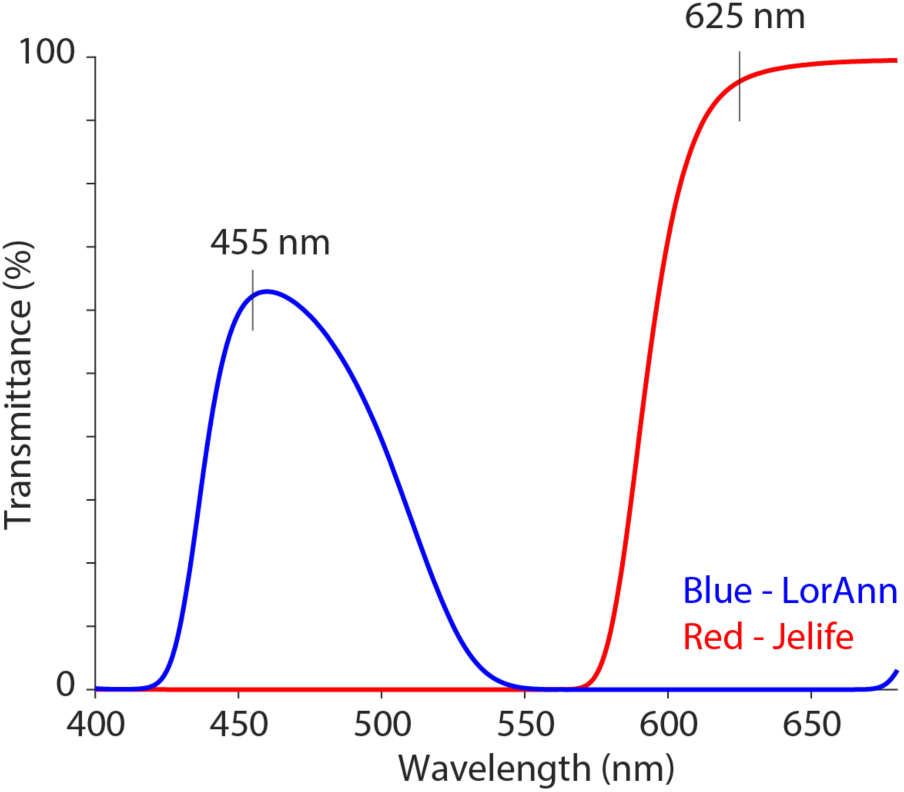
We measured transmittance for the red (Jelife Red) and blue dyes (LorAnn Blue) that are used to paint the right and left antennae, respectively. Note the clean separation between red and blue dye curves, around the blue (455nm) and red (625nm) LEDs chosen for this experiment.

**Fig. S3.**
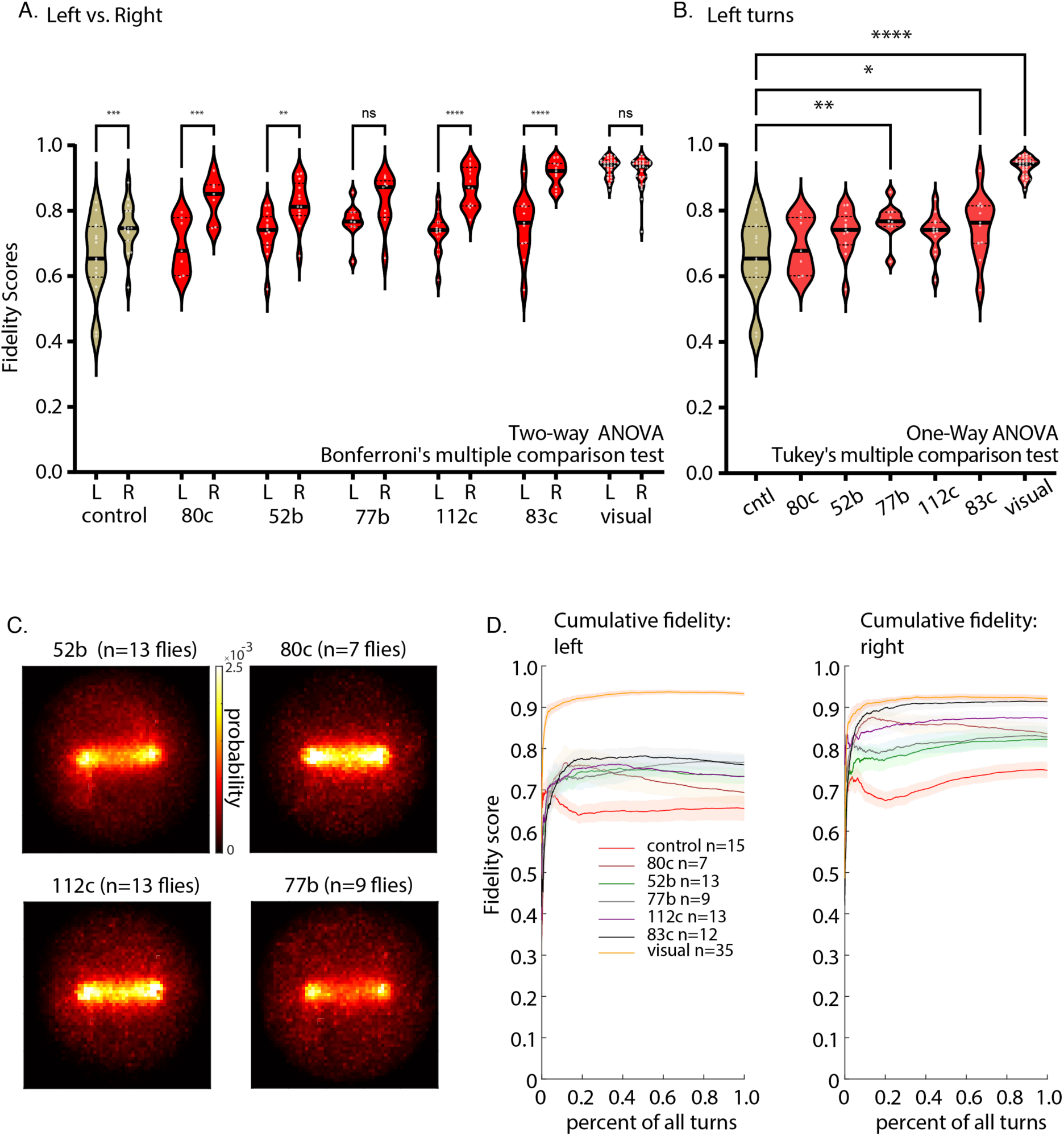
A. Fidelity scores for left vs. right turns, for each MBON line chosen for this experiment. Overall, red light stimulation, leading to right turning is more effective at following the guidance cue than left turning. Two way ANOVA with Bonferroni multiple comparison correction was used. B. Fidelity scores for left turns. One way ANOVA with Tukey multiple comparison correction was used. C. Probability spatial histograms for MBON lines that were not shown in Figure 3. Overall all MBON lines improved performance of the A to B guidance task over the control. D. Cumulative fidelity scores for blue light driven left turns, showing a modest improvement over controls and red light driven right turns. Particularly the red light stimulation of the line 83c is comparable to visual stimulation.

**Fig. S4.**
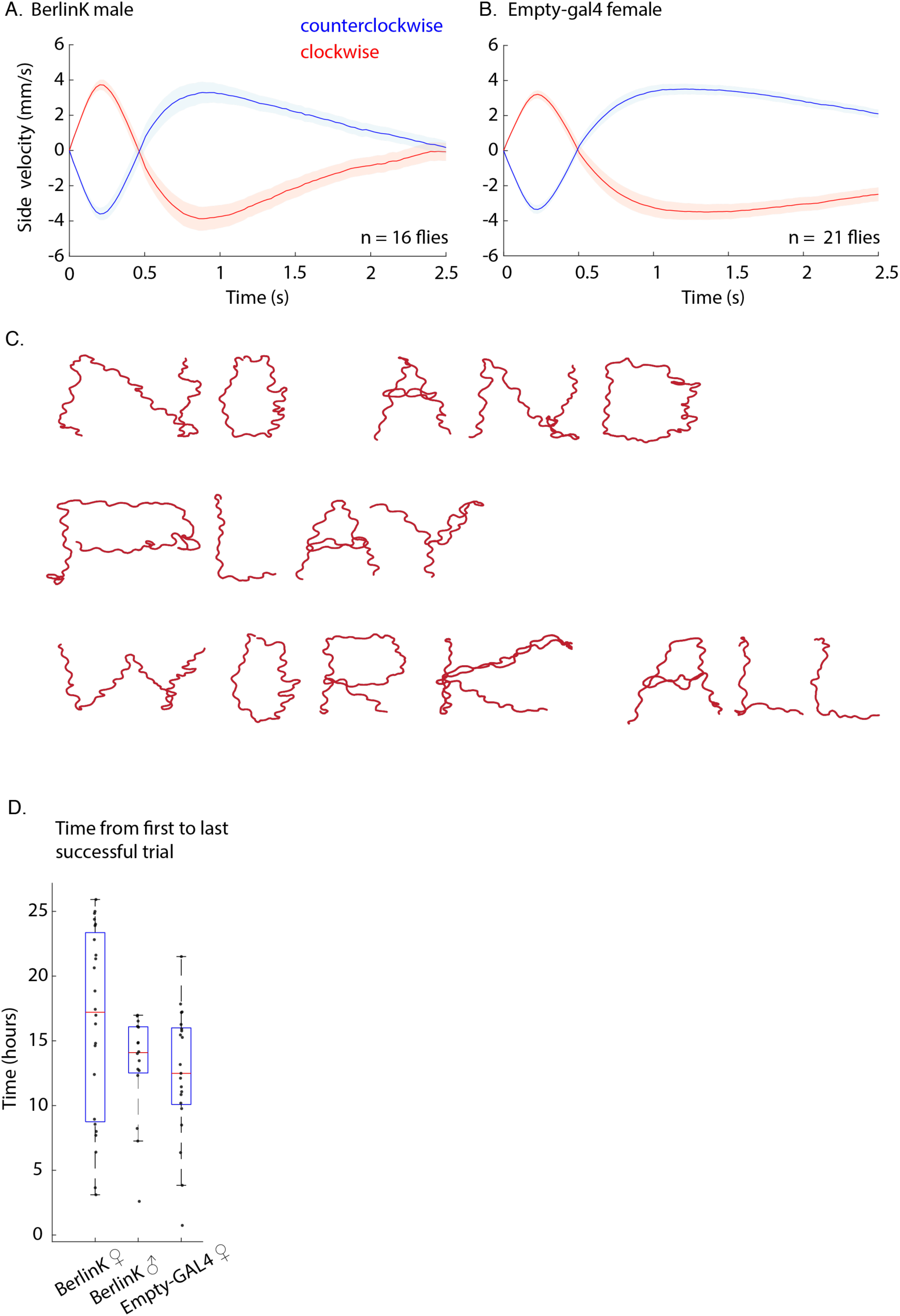
Similar A) BerlinK males and B) Empty-GAL4 female side velocity profiles to clockwise and counterclockwise rotation of the pinwheel. Time zero is the onset of rotation. C. Additional example of text writing by a single fly. D. Time between first and last successful trial for each genotype, illustrating that flies can perform this task for hours. Number of flies is same as Figure 1G.

**Fig. S5.**
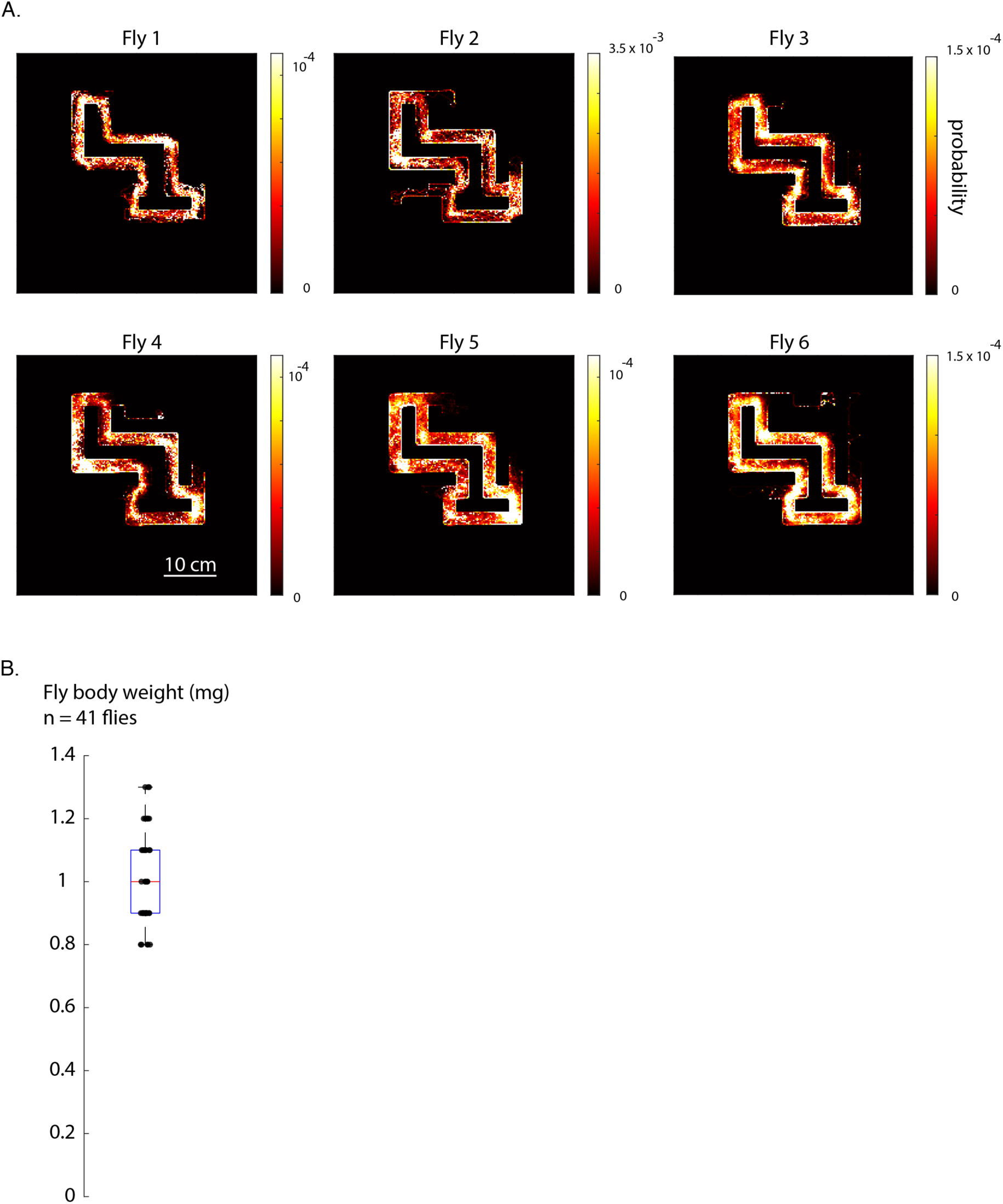
A. Probability spatial heatmaps for each fly visually guided through the “main route” of the maze. B. Body weight of flies that were used in weight carrying experiments (n = 41 flies).

**Fig. S6.**
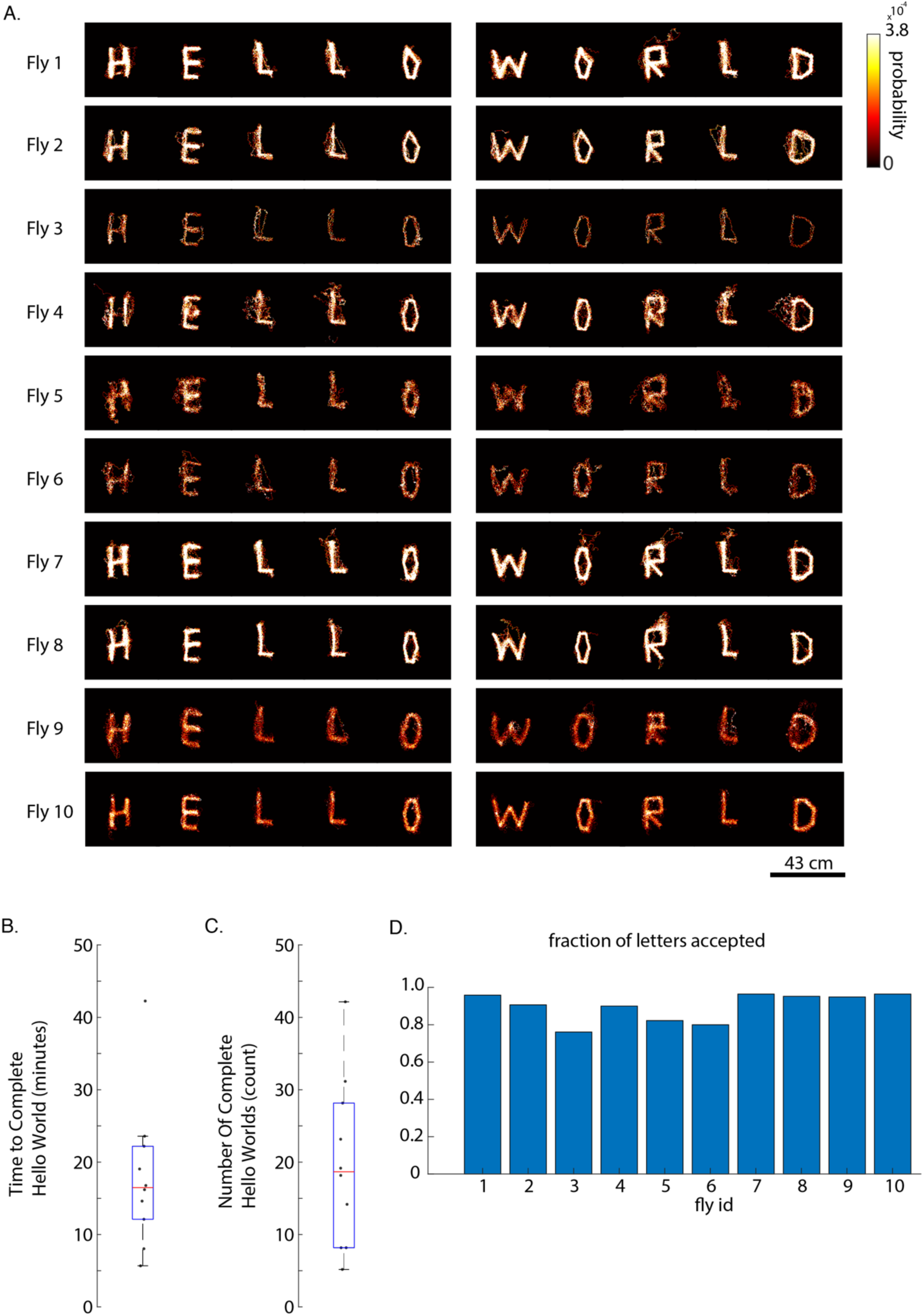
A. Probability spatial histograms for all 11 flies in our dataset that were visually guided to ‘write’ HELLO WORLD. 3 of these flies are shown in Figure 4. Visual guidance in this task is consistent across flies. B. Each data point is the median time to complete all the letter patterns of ‘Hello World’ for each fly (n = 10 flies). C. Each data point is the number of completed ‘Hello World’ letter patterns for each fly (n = 10 flies). D. Fraction of letters, from all attempted letters, that are shown in A. A letter was disqualified from display if there were more than 10 failed attempts to a goal for any of the goals or if the path to any of the goals in a letter deviated by more than 12.5 cm.

**Fig. S7.**
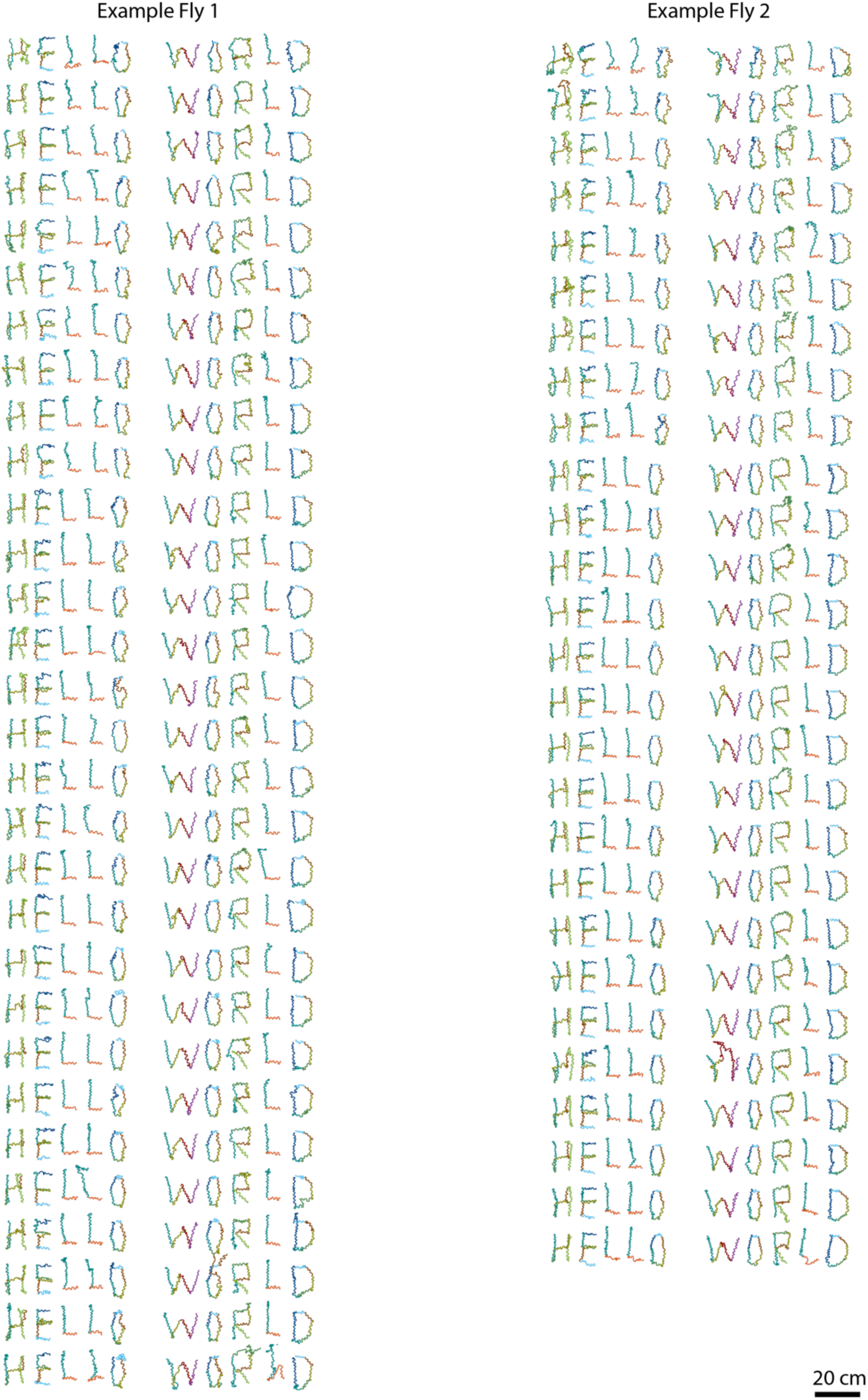
Individual HELLO WORLD instances from fly 7 (fly 1) and 8 (fly 2) in Supplementary Figure 6.

**Table S1.** Genotypes of flies described in each figure.

| **Figure** | **Genotypes** |
| --- | --- |
| 1C-1E | *Berlin-K* wild type strain M (Stock# 8522, BDSC) |
| 1F-1G, 1H | *Berlin-K* wild type strain M & *empty-Gal4* |
| 2B-2H | *UAS-CsChrimson; Orco-Gal4; UAS-ChR2/\*empty-Gal4*<br>*\*empty-Gal4* = *w[1118]; P(y[+t7.7] w[+mC]=GAL4.1Uw)attP2* (Stock# 68384, BDSC) |
| 3C-3E | *UAS-CsChrimson; Orco-Gal4; UAS-ChR2/empty-Gal4* (control)<br>*UAS-CsChrimson; Orco-Gal4; UAS-ChR2/R50A05-GAL4.DBD, R33E02-p65.AD* (MB080C)<br>*UAS-CsChrimson; Orco-Gal4/R71D08-p65ADZp; UAS-ChR2/R11F03-ZpGdbd* (MB052B)<br>*UAS-CsChrimson; Orco-Gal4/R25D01-p65.AD; UAS-ChR2/R19F09-GAL4.DBD* (MB077B)<br>*UAS-CsChrimson; Orco-Gal4; UAS-ChR2/R13F04-GAL4.DBD, R93D10-p65.AD* (MB112C)<br>*UAS-CsChrimson; Orco-Gal4; UAS-ChR2/R94B10-GAL4.DBD, R52G04-p65.AD* (MB083C)<br>*Berlin-K* wild type strain M (visual) |
| 3F | *UAS-CsChrimson; Orco-Gal4; UAS-ChR2/empty-Gal4* (control)<br>*UAS-CsChrimson; Orco-Gal4; UAS-ChR2/R94B10-GAL4.DBD, R52G04-p65.AD* (MB083C) |
| 4B-4C | *Berlin-K* wild type strain M |
| 4D | *UAS-CsChrimson; Orco-Gal4; UAS-ChR2/empty-Gal4* &<br>*UAS-CsChrimson; Orco-Gal4; UAS-ChR2/R50A05-GAL4.DBD, R33E02-p65.AD* |
| 5A-5B | *Berlin-K* wild type strain M |
| 6B-6F | *Berlin-K* wild type strain M |
| 6G-6K | *empty-Gal4* |

**Table S2.**
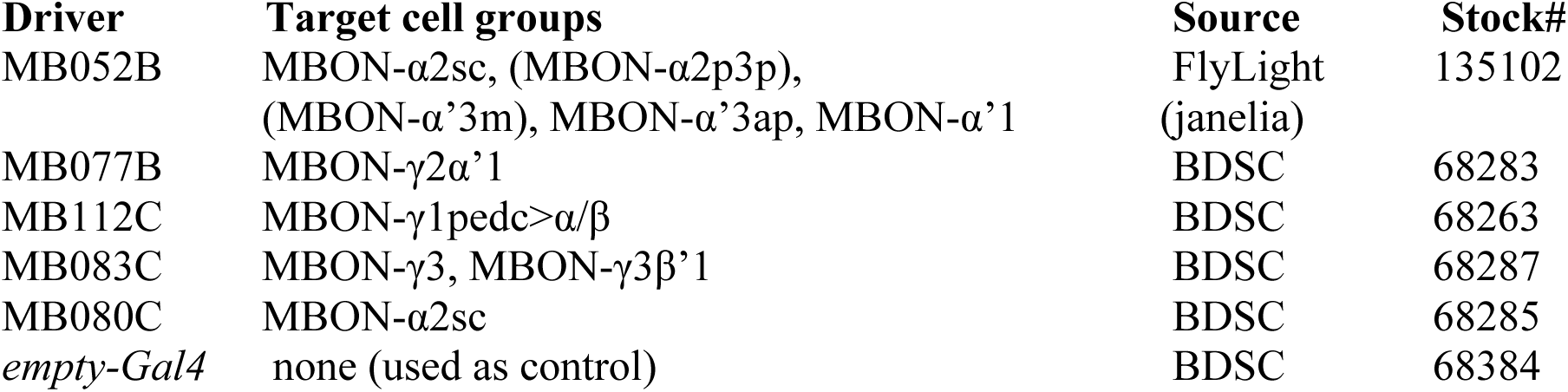
Fly strains tested for the MBON screen.

| **Driver** | **Target cell groups** | **Source** | **Stock#** |
| --- | --- | --- | --- |
| MB052B | MBON-α2sc, (MBON-α2p3p),<br>(MBON-α'3m), MBON-α'3ap, MBON-α'1 | FlyLight<br>(janelia) | 135102 |
| MB077B | MBON-γ2α'1 | BDSC | 68283 |
| MB112C | MBON-γ1pedc>$\alpha$/$\beta$ | BDSC | 68263 |
| MB083C | MBON-γ3, MBON-γ3β'1 | BDSC | 68287 |
| MB080C | MBON-α2sc | BDSC | 68285 |
| *empty-Gal4* | none (used as control) | BDSC | 68384 |

**Table S3:**
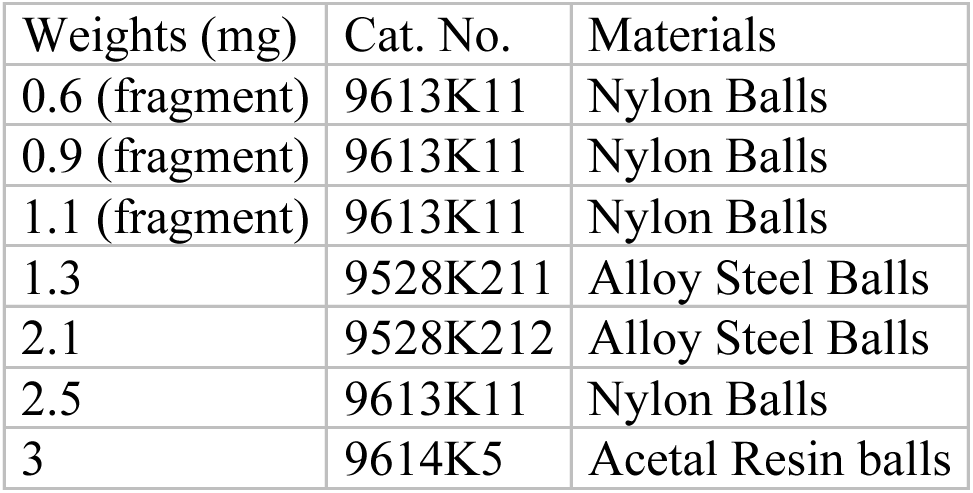
Ball information (Manufacturer: McMaster-Carr)

| Weights (mg) | Cat. No. | Materials |
| --- | --- | --- |
| 0.6 (fragment) | 9613K11 | Nylon Balls |
| 0.9 (fragment) | 9613K11 | Nylon Balls |
| 1.1 (fragment) | 9613K11 | Nylon Balls |
| 1.3 | 9528K211 | Alloy Steel Balls |
| 2.1 | 9528K212 | Alloy Steel Balls |
| 2.5 | 9613K11 | Nylon Balls |
| 3 | 9614K5 | Acetal Resin balls |

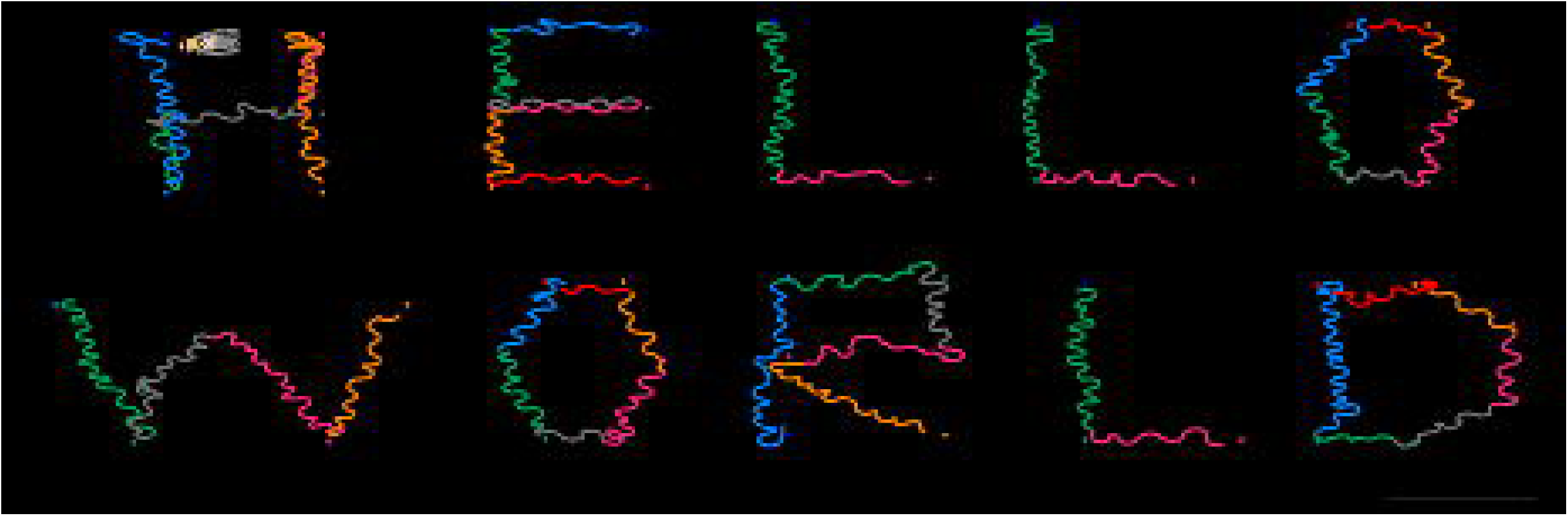

